# Extrinsic and Intrinsic Dynamics in Movement Intermittency

**DOI:** 10.1101/402552

**Authors:** Damar Susilaradeya, Wei Xu, Thomas M Hall, Ferran Galán, Kai Alter, Andrew Jackson

**Affiliations:** Institute of Neuroscience, Faculty of Medical Sciences, Newcastle University, Newcastle NE2 4HH, UK; Current address: Department of Basic Neuroscience, Faculty of Medicine, University of Geneva, CH-1202 Genève

**Keywords:** Submovements, movement intermittency, motor cortex, optimal feedback control

## Abstract

What determines how we move in the world? Motor neuroscience often focusses either on intrinsic rhythmical properties of motor circuits or extrinsic sensorimotor feedback loops. Here we show that the interplay of both intrinsic and extrinsic dynamics is required to explain the intermittency observed in continuous tracking movements. Using spatiotemporal perturbations in humans, we demonstrate that apparently discrete submovements made 2-3 times per second reflect constructive interference between motor errors and continuous feedback corrections that are filtered by intrinsic circuitry in the motor system. Local field potentials in monkey motor cortex revealed characteristic signatures of a Kalman filter giving rise to both low-frequency cortical cycles during movement, and delta oscillations during sleep. We interpret these results within the framework of optimal feedback control, and suggest that the intrinsic rhythmicity of motor cortical networks reflects an internal model of external dynamics which is used for state estimation during feedback-guided movement.

## Introduction

Many visually-guided movements are characterized by intermittent speed fluctuations. For example while tracking slowly-moving targets, humans make around 2-3 submovements per second. Although first described over a century ago (Woodworth, 1899; Craik, 1947; Vince, 1948) the cause of movement intermittency remains debated. Submovements often disappear in the absence of vision (Miall *et al.,* 1993a) and are influenced by feedback delays (Miall, 1996), suggesting their timing depends on extrinsic properties of visuomotor feedback loops. However, rhythmicity is also reported in the absence of feedback (Doeringer and Hogan, 1998), and it has been suggested that an internal refractory period, clock or oscillator parses complex movements into discrete isochronal segments (Viviani and Flash, 1995; Russell and Sternad, 2001; Loram *et al.,* 2006; Hogan and Sternad, 2012). Cyclical dynamics within motor cortical networks with a time period of 300-500ms may reflect the neural correlates of such an intrinsic oscillator (Churchland *et al.,* 2012; Hall *et al.,* 2014). During continuous tracking, each submovement is phase-locked to a single cortical cycle, giving rise to low-frequency coherence between cortical oscillations and movement speed (Jerbi *et al.,* 2007; Hall *et al.,* 2014; Pereira *et al.,* 2017). Moreover, this rhythmicity appears conserved across a wide range of behaviors and even shares a common dynamical structure with delta oscillations during sleep (Hall *et al.,* 2014). It has been proposed that recurrent networks express these intrinsic dynamics as an ‘engine of movement’ responsible for internal generation and timing of the descending motor command (Churchland *et al.,* 2012). Nevertheless, the interplay between intrinsic rhythmicity and extrinsic feedback remains poorly understood. For example, if feedback delays influence submovement timing they might be expected also to alter the frequency of cortical cycles. However, this seems incompatible with conserved intrinsic dynamics evident across multiple behavioral contexts including sleep. Moreover, the precise computational role of such intrinsic circuitry remains uncertain.

In recent years, stochastic optimal control theory has emerged as an influential computational approach to understanding human movement, due to recognition of the impact of noise in both motor and sensory signals on behavior (Todorov and Jordan, 2002; Scott, 2004). In the presence of delayed, uncertain measurements, feedback should act on optimal estimates of the discrepancy between desired goals and current motor states. Optimal feedback control (OFC) explains many features of movement but it is unclear whether optimality principles alone can account for movement intermittency. Various modifications to OFC have been proposed, for example explicitly including a refractory period between submovements (Gawthrop *et al.,* 2011), but theoretical justification for these additions is lacking. Here we present evidence from visuomotor tracking by humans and non-human primates in support of an OFC-based model of movement intermittency that does not require explicit parsing of submovements. Instead, continuous integration of external feedback with internal state estimation provides a framework for understanding both extrinsic and intrinsic contributions to intermittency. This can account for many puzzling features of submovements, and provides a parsimonious explanation for conserved cyclical dynamics in motor cortex networks during behavior and sleep.

## Results

### Submovements reflect constructive interference between motor noise and delayed feedback corrections

Human subjects generated bimanual isometric index finger forces to track targets that moved in 2D circular trajectories with constant speed (Fig. 1A). We measured intermittency in the angular velocity of the cursor (Fig. 1B C), using spectral analysis to quantify submovement frequencies. Under normal feedback conditions, power spectra generally exhibited a principal peak at around 2 Hz (Fig. 1D) and this frequency was only slightly affected by target speed (Figure S1), consistent with previous descriptions of movement intermittency (Miall, 1996). However, submovement frequencies were markedly altered when visual feedback of the cursor was delayed relative to finger forces. With delays of 100 and 200 ms, the frequency of the primary peak reduced to around 1.4 and 1 Hz respectively (Fig. 1D, Fig. S1, Fig. S2), suggesting submovement timing was not determined by a fixed internal clock but depended instead on extrinsic feedback properties. Interestingly, a further peak appeared at approximately three times the frequency of the primary peak and, with increased delays of 300 and 400 ms, a 5^th^ harmonic was observed. The time-periods of the first, third and fifth harmonics were linearly related to extrinsic delay times with gradients of 1.89 ± 0.20, 0.59 ± 0.04 and 0.33 ± 0.11 respectively (Fig. 1E, Table S1).

**Figure 1.**
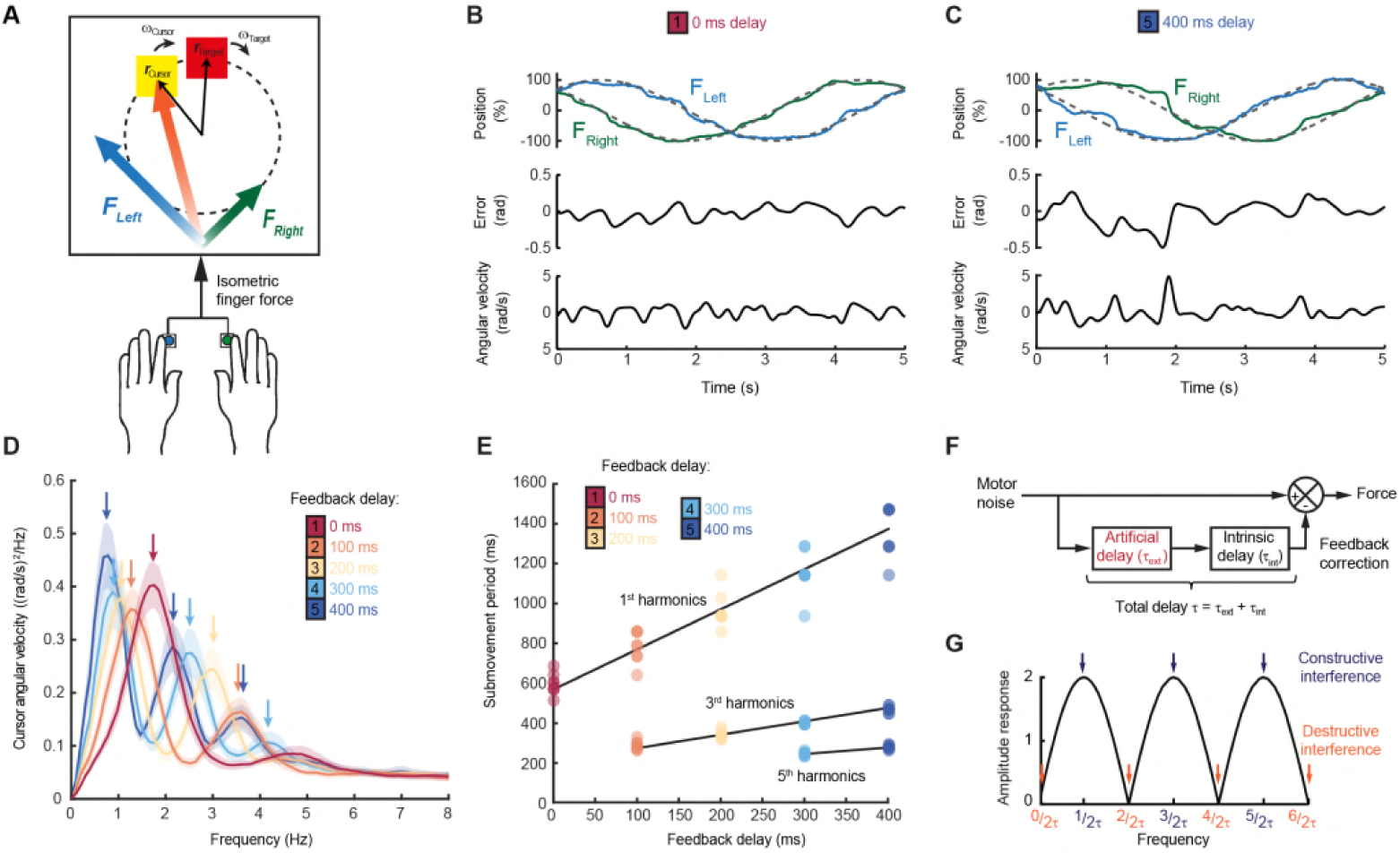
Movement intermittency during visuomotor tracking depends on feedback delays. (A) Schematic of human tracking task. Bimanual isometric finger forces control 2D cursor position to track slow, circular target motion. Kinematic analyses use the angular velocity of the cursor subtended at the screen center screen. (B) Example force (*top*), angular error (*middle*) and cursor angular velocity (*bottom*) traces during target tracking with no feedback delay. Submovements are evident as intermittent fluctuations in angular velocity. (C) Example movement traces with 400 ms feedback delay. (D) Power spectra of cursor angular velocity with different feedback delays between 0–400 ms. Average of 8 subjects, shading indicates standard error of mean (s.e.m.). See also Figure S2. (E) Submovement periods (reciprocal of the peak frequency for each harmonic) for all subjects with different feedback delays. See also Table S1. (F) Schematic of a simple delayed feedback controller. (G) Amplitude response of the system shown in (F), known as a comb filter.

These results are consistent with a feedback controller responding to broad-spectrum (stochastic) tracking errors introduced by noise in the motor output, for which the response is delayed by time *τ* (Fig. 1F). In signal processing terms, subtracting a delayed version from the original signal is known as comb filtering. For motor noise components with a time period, 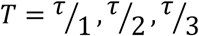 …, delayed feedback accurately reflects current errors, resulting in regularly spaced notches in the amplitude response of the system (Fig. 1G) and attenuation from the resultant cursor movement through destructive interference. By contrast, for motor noise with a time-period, 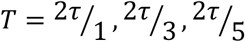 …, delayed feedback is exactly out-of-phase with the current error. Thus, corrective movements exacerbate these components through constructive interference leading to spectral peaks at frequencies:

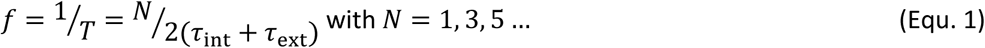

Submovement frequencies in our data approximately matched this model assuming the total feedback delay comprised the experimental manipulation *τ*_ext_ added to a constant physiological response latency *τ*_int_ of around 300 ms (Table S1), comparable to visual reaction times.

Note that in this interpretation, intermittency arises not from active generation of discrete submovement events but as a byproduct of continuous, linear feedback control with inherent time delays. Submovement frequencies need not be present in the smooth target movement, nor do they arise from controller non-linearities. Instead these frequencies reflect components of broad-band motor noise that are exacerbated by delayed feedback corrections. To seek further evidence that intermittency arises from constructive interference between motor noise and delayed feedback corrections, we generated artificial errors during target tracking by adding spatial perturbations to the cursor displayed to subjects. Within individual trials, a sinusoidal displacement was applied in a direction aligned to target motion and at a frequency between 1-5 Hz. Perturbation amplitudes were scaled to have equivalent peak angular velocities (equal to the angular velocity of the target). Our hypothesis was that artificial errors at submovement frequencies would be harder to track (because of constructive interference) than perturbations at frequencies absent from the velocity spectrum.

**Figure 1—source data 1.** This spreadsheet contains the frequencies of spectral peaks and associated regression analysis shown in Figure 1D,E. These data can be opened with Microsoft Excel or with open-source alternatives such as OpenOffice.

Figure 2A shows example tracking behavior with a 2 Hz perturbation. Note that the peak angular velocity of force responses (*black line,* calculated from the subject’s finger forces) occurred around the same time as the peak angular velocity of the perturbation (*green line*). As a result, the angular velocity of the cursor (*yellow line,* reflecting the combination of the subject’s forces with the perturbation) exhibited pronounced oscillations that were larger than the perturbation. Figure 2B shows performance in the same task when visual feedback was delayed by 200 ms. In this condition, peaks in force velocity coincided with perturbation troughs, attenuating the disturbance to cursor velocity. Figure 2C,D and Figure S3 overlay cursor velocity spectra in the presence of each perturbation frequency (with feedback delays of 0 and 200 ms). As previously, in the absence of feedback delay the frequency of submovements was around 2 Hz. Correspondingly, perturbations at 2 Hz induced a large peak in the cursor velocity spectrum, indicating that the artificial error was not effectively tracked. By contrast, with a feedback delay of 200 ms the cursor velocity spectrum with a 2 Hz perturbation was attenuated. The largest spectral peaks were instead associated with 1 and 3 Hz perturbations, matching the frequencies of submovements in this delay condition. Figure 2E shows the amplitude response of cursor movements (the relative amplitude of cursor movements phase-locked to the perturbation) at each frequency for both delay conditions. Cursor amplitude responses greater than unity at 2 Hz (with no delay), and at 1 and 3 Hz (with 200 ms delay) indicate exacerbation of intermittencies introduced by artificial errors at submovement frequencies. Analysis of variance (ANOVA) with two factors (delay time and perturbation frequency) revealed a highly significant interaction (n=8 subjects, F_4,70_=110.2, P<0.0001), confirming the interdependence of feedback delays and frequencies of constructive/destructive interference.

**Figure 2.**
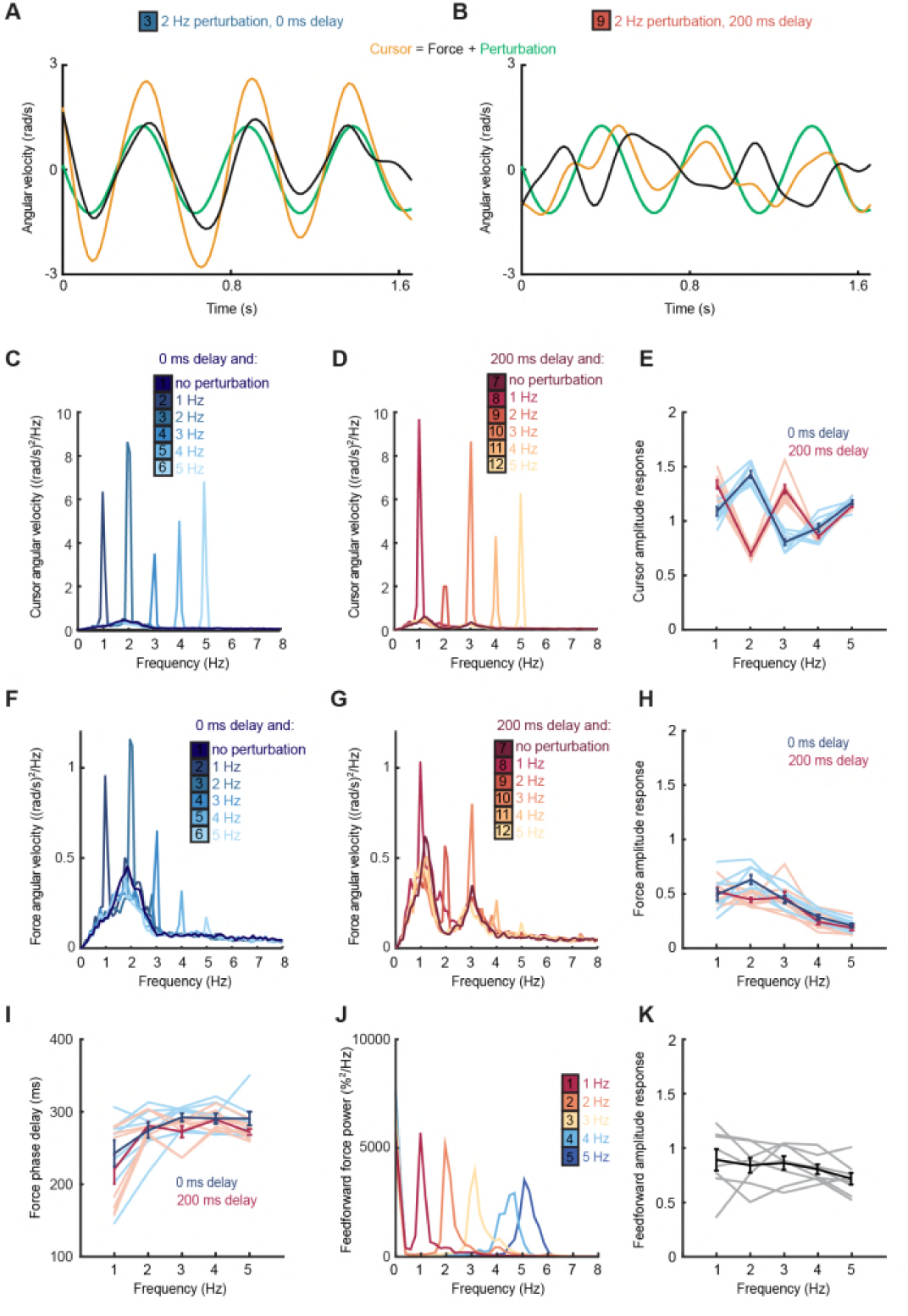
Frequency responses and phase delays to artificial motor errors. (A) Example force (*black*) and cursor (*yellow*) angular velocity traces in the presence of a 2 Hz perturbation (*green*). No feedback delay is added. The force response and perturbation sum to produce large fluctuations in cursor velocity. (B) Comparable data with a feedback delay of 200 ms. In this condition, force responses cancel the perturbation leading to an attenuation of intermittency. (C) Power spectra of cursor angular velocity with 1–5 Hz perturbations and no feedback delay. Average of 8 subjects. See also Figure S3. (D)Power spectra of cursor angular velocity with 1–5 Hz perturbations and 200 ms feedback delay. (E) Cursor amplitude response to 1–5 Hz perturbations with no feedback delay (*blue*) and 200 ms feedback delay (*red*) for individual subjects. Also shown is average ± s.e.m. of 8 subjects. (F) Power spectra of force angular velocity with 1–5 Hz perturbations and no feedback delay. See also Figure S4. (G) Power spectra of force angular velocity with 1–5 Hz perturbations and 200 ms feedback delay. (H) Force amplitude response to 1–5 Hz perturbations with no feedback delay (*blue*) and 200 ms feedback delay (*red*). Also shown is average ± s.e.m. of 8 subjects. (I) Intrinsic phase delay of force response to 1–5 Hz perturbations with no feedback delay (*blue*) and 200 ms feedback delay (*red*). Also shown is average ± s.e.m. of 8 subjects. (J) Power spectrum of finger forces generated in the feedforward task with auditory cues at 1-5 Hz. Average of 8 subjects. See also Figure S5. (K) Force amplitude response to auditory cues in the feedforward task. Also shown is average ± s.e.m. of 8 subjects.

### Feedback responses reflect filtered visual discrepancies

It is clear from the velocity spectra in Figure 1D that not all submovement harmonics predicted by the comb filter model were present with the same amplitude within our tracking data. Rather, intermittency peaks for each delay condition were embedded within a broad low-pass envelope. Next we considered the origin of this delay-independent envelope. Our first hypothesis was that this might reflect the spectral content of motor noise during tracking. However we could reject this as the sole contributing factor by examining the force amplitude response to perturbations (the relative amplitude of force responses phase-locked to the perturbation). Figures 2F,G and Figure S4 show power spectra of the angular velocity derived from subject’s forces, under feedback delays of 0 and 200 ms. Figure 2H shows the corresponding force amplitude response at each frequency. Analyzed in this way, amplitude responses were largely independent of extrinsic delay. However, as with submovement peaks, feedback responses were also attenuated at higher frequencies. A two-factor ANOVA confirmed a significant main effect of frequency (n=8 subjects, F_4,70_=36.3, P<0.0001) but not delay time (F_1,70_=3.1, P=0.08), and only a weakly significant interaction (F_4,70_=2.9, P=0.03). Moreover, the phase delay of force responses was reduced at low frequencies (Fig. 2I). As with the amplitude response, there was a significant effect of frequency (F_4,70_=9.5, P<0.0001) but not extrinsic delay (F_1,70_ =2.6, P=0.12) on this phase delay, and no significant interaction (F_4,70_=0.7, P=0.6). In other words, feedback corrections to artificial noise with equal amplitude at different frequencies revealed characteristic signatures of a filter that was independent of extrinsic feedback delays. Moreover, this intrinsic filter had the appropriate bandwidth to account for attenuation of intermittency at higher frequencies.

**Figure 2—source data 2.** This spreadsheet contains the cursor/force/feedforward amplitude response and phase delay data shown in Figure 2E,H,I,K. These data can be opened with Microsoft Excel or with open-source alternatives such as OpenOffice.

Next we considered whether this attenuation was a property of motor pathways, for example reflecting filtering by the musculoskeletal system. However, it is well-known that the frequencies of feedforward movements can readily exceed submovement frequencies observed during feedback-guided behavior (Kunesch *et al.,* 1989). We confirmed this by asking subjects to produce force fluctuations of a defined amplitude, but without providing a moving target to track. Instead we used auditory cues (a metronome) to indicate the required movement frequency. In this case, subjects could generate force fluctuations up to 5 Hz with little attenuation (Fig. 2J,K and Fig. S5). Therefore we concluded that filtering during visuomotor tracking was not inherent to the motor pathway and considered instead whether it could be a property of the feedback loop.

### Filtered feedback corrections are consistent with optimal state estimation

The visual system can perceive relatively high frequencies (up to flicker-fusion frequencies above 10 Hz). However, for movements in the physical world, it is unlikely that high-frequency tracking discrepancies reflect genuine motor errors, since this would imply implausibly large accelerations of the body. Given inherent uncertainties in sensation, an optimal state estimator should attribute such errors to sensory noise (as this is unconstrained by Newtonian dynamics). Formally, the task of distinguishing the true state of the world from uncertain, delayed measurements can be achieved by a Kalman filter which continuously integrates new evidence with updated estimates of the current state evolving according to a model of the external dynamics (Fig. 3A). For simplicity we assumed the 1D position of the body (cursor) should move with constant velocity relative to the slow, predictable target unless acted upon by accelerative forces, leading to a two-dimensional state transition model:

**Figure 3.**
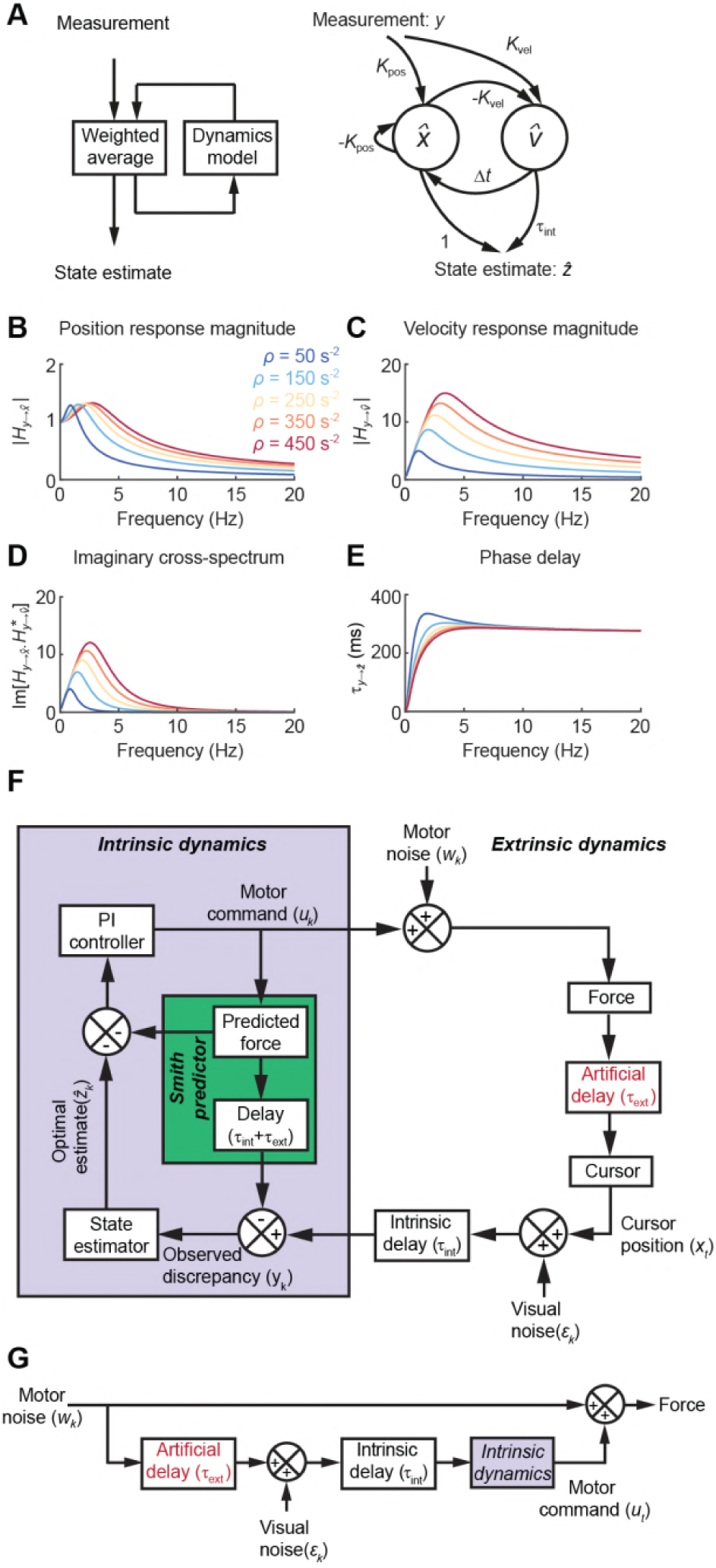
State estimation with a Kalman filter. (A) *Left:* Schematic of a Kalman filter. Noisy measurements are combined with an internal model of the external dynamics to update an optimal estimate of current state. *Right:* A dynamical system for optimal estimation of position, based on an internal model of position and velocity. (B, C) Magnitude response of transfer function from measurement to position and velocity estimates for the Kalman filter with different ratios of process to measurement noise (*ρ*). (D) Imaginary component of cross-spectrum between position and velocity transfer functions. (E) Phase delay of optimal estimate of position based on delayed measurement of position. (F) Schematic of optimal feedback controller model incorporating state estimation and a Smith Predictor architecture to accommodate feedback delays. (G) Simplified rearrangement of (F), showing the feedforward relationship between motor noise and force output. This rearrangement is possible because the Smith Predictor prevents motor corrections reverberating multiple times around the feedback loop.

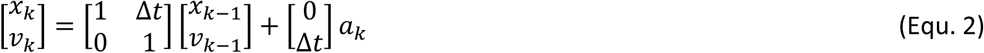

where *x*_*k*_ and *v*_*k*_ are the relative position and velocity of the cursor at time-step *k,* Δ*t* is the interval between time-steps, and the process noise 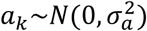. Visual feedback, *y*_*k*_, was assumed to comprise a noisy measurement of relative position:

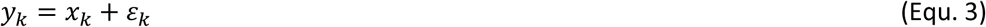

with measurement noise 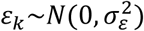. Optimal estimates of relative position and velocity, 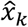 and 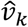 are given by a steady-state Kalman filter of the form:

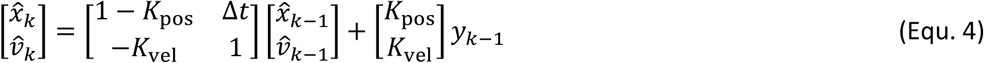

The innovation gains *K*_pos_ and *K*_vel_ depend only on the ratio of process to measurement noise, 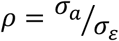, which in turn determines the cut-off frequency above which measurements are filtered 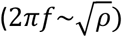. Figure 3B,C shows the amplitude response for position and velocity estimates. Note that these are out of phase with each other, and therefore broadband input results in a complex cross-spectral density between them. The imaginary component of this cross-spectrum exhibits a characteristic resonance peak (Fig. 3D). Feedback delays can be accommodated by projecting the state estimate forward in time:

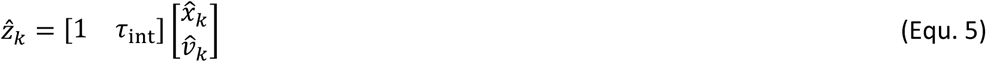

The phase delay of the optimal position estimate for the current state, 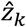, falls towards zero at low frequencies, consistent with successful prediction on the basis of delayed measurement (Fig. 3E). This steady-state Kalman filter was incorporated within a 1D feedback controller (Fig. 3F; see Methods for details) which also included an internal feedback loop to cancel the sensory consequences of motor commands. This architecture, known as a Smith Predictor (Miall *et al.,* 1993b; Abe and Yamanaka, 2003), prevents corrections from reverberating around the external feedback loop, such that the resultant closed-loop behavior is formally equivalent to the simpler feedforward system shown in Figure 3G. This rearrangement provides a useful intuition about our behavioral results. Tracking errors (due to motor noise) drive feedback corrections that are delayed, corrupted (by sensory noise) and filtered (by intrinsic dynamics). The power spectrum of the resultant movement reflects constructive/destructive interference between feedback corrections and the original tracking error.

This simple model readily accounted for the main features of our human data, including the cursor amplitude response to perturbations (Fig. 4A-E), and the low-pass filtering (Fig. 4F-H) and phase delay (Fig. 4I) of force responses. Moreover, because of frequency-dependent phase delays introduced by state estimation, the model predicted that precise frequencies of submovement peaks should deviate slightly from those calculated using a constant physiological response latency. This effect was confirmed in our behavioral data by calculating (with Equ. 1) the intrinsic delay time corresponding to each spectral harmonic under all feedback delays (Fig. 1D). This intrinsic delay time was positively correlated with the frequency of the harmonic (n=11 spectral peaks, R=0.85, P=0.0009; Fig. 4J). Finally, overall tracking performance (as measured by the root mean squared positional error over time) matched well with subjects’ actual performance across conditions (Fig. 4K). Note that irrespective of delay, the lowest frequency perturbation was associated with the greatest positional error (since perturbations had equal peak-to-peak velocity and were therefore larger in amplitude at low frequencies). However, performance was most affected by the 1 Hz perturbation with a 200 ms delay, corresponding to a frequency of constructive interference.

**Figure 4.**
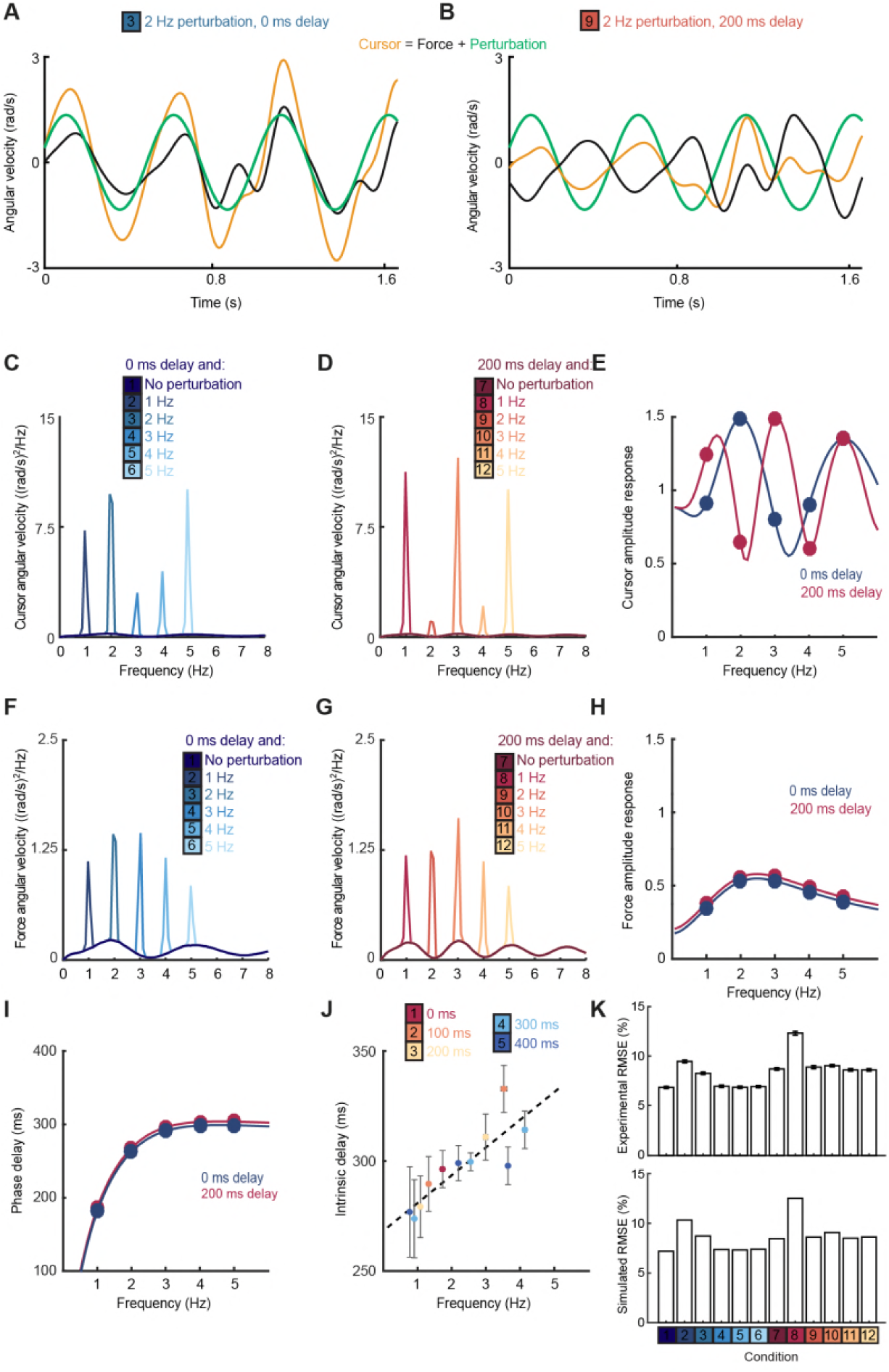
OFC model reproduces human behavioral data. (A) Simulated tracking performance of the OFC model with a 2 Hz sinusoidal perturbation and no feedback delay. (B) Simulated tracking performance of the OFC model with a 2 Hz sinusoidal perturbation and 200 ms feedback delay. (C) Power spectrum of simulated cursor velocity with 1–5 Hz perturbations and no feedback delay. (D) Power spectrum of simulated cursor velocity with 1–5 Hz perturbations and 200 ms feedback delay. (E) Simulated cursor amplitude response to 1–5 Hz perturbations with no feedback delay (*blue*) and 200 ms feedback delay (*red*). (F) Power spectrum of simulated force velocity with 1–5 Hz perturbations and no feedback delay. (G) Power spectrum of simulated force velocity with 1-5 Hz perturbations and 200 ms feedback delay. (H) Simulated force amplitude response to 1–5 Hz perturbations with no feedback delay (*blue*) and 200 ms feedback delay (*red*). (I) Simulated intrinsic phase delay of force responses to 1–5 Hz perturbations with no feedback delay (*blue*) and 200 ms feedback delay (*red*). (J) Intrinsic delay times corresponding to all submovement peaks/harmonics in Figure 1D, plotted against the frequency of the peak. Error bars indicate s.e.m. across 8 subjects. (K) *Top:* Positional inaccuracy of human tracking for all conditions quantified as root mean squared error (RMSE). Average ± s.e.m. of 8 subjects. *Bottom:* RMSE of simulated tracking for all conditions.

In summary, amplitude and phase responses to perturbations during human visuomotor tracking provide compelling evidence for intrinsic filtering of measurement noise from feedback corrections, while a plausible computational justification is provided by optimal state estimation. Moreover, while this interpretation is derived from computational principles, the schematic on the right of Fig. 3A suggests how a steady-state Kalman filter could be implemented by neural circuitry. Two neural populations representing position and velocity should evolve according to Equ. 4 and thus exhibit a resonance peak in their imaginary cross-spectrum. To seek further evidence for the neural implementation of such a filter we turned to intracortical recordings in non-human primates.

### Movement intermittency in a non-human primate tracking task

We were interested in whether cyclical motor cortex dynamics could reflect the neural correlates of the two interacting neural populations described above, and thereby account for filtering of feedback responses during visuomotor tracking. We therefore analyzed local field potential (LFP) recordings from monkey primary motor cortex (M1) during a center-out isometric wrist torque task that we have used previously to characterize both submovement kinematics and population dynamics (Hall *et al.,* 2014). Figure 5 shows example tracking behavior (Fig. 5A), radial cursor velocity (Fig. 5B) and multichannel LFPs (Fig. 5C) as monkeys moved to peripheral targets under two feedback delay conditions. Movement intermittency was apparent as regular submovement peaks in the radial cursor velocity. Moreover LFPs exhibited low-frequency oscillations during movement, with a variety of phase-shifts present on different channels. Principal component analysis (PCA) yielded two orthogonal components of the cortical cycle (Fig. 5E), and the close coupling with submovements was revealed by overlaying the cursor velocity profile onto, in this case, the second principal component (PC) (Fig. 5E).

**Figure 5.**
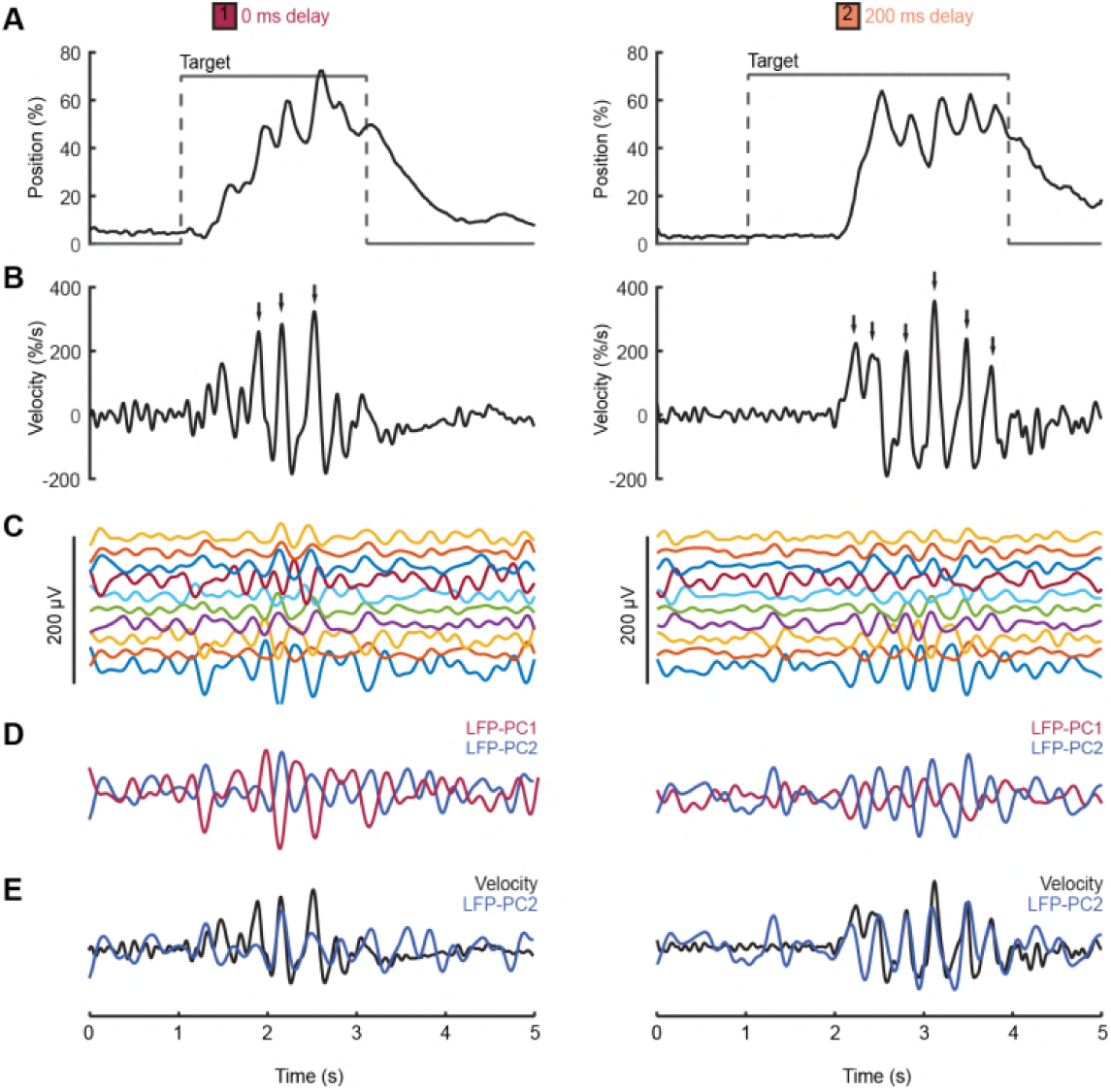
Movement intermittency in a non-human primate tracking task. (A) Radial cursor position during a typical trial of the center-out isometric wrist torque task under two different feedback delay conditions. Data from Monkey U. (B) Radial cursor velocity. Arrowheads indicate time of submovements identified as positive peaks in radial cursor velocity >150%/s. (C) Low-pass filtered, mean-subtracted LFPs from M1. (D) First two principal components (PCs) of the LFP. (E) The second LFP-PC overlaid on the radial cursor velocity.

As with humans, in the absence of feedback delay the cursor velocity (after removing task-locked components, see Methods) was dominated by a single spectral peak around 2-3 Hz (Fig. 6A,E; *top red traces*). A broad peak at approximately the same frequency was also observed in average LFP power spectra (Fig. 6B,F), while coherence analysis confirmed consistent phase-coupling between LFPs and cursor velocity (Fig. 6C,G). We also calculated imaginary coherence spectra between pairs of LFPs (see Methods) to separate local signal components with a consistent, non-zero phase difference from in-phase components (e.g. due to volume conduction from distant sources), which revealed more clearly the 2-3 Hz LFP oscillation (Fig. 6D,H). An obvious interpretation of these results could be that oscillatory activity in the motor system drives submovements in a feedforward manner. In this case we would expect the frequency of the cortical oscillation to reliably reflect the intermittency observed in behavior.

**Figure 6.**
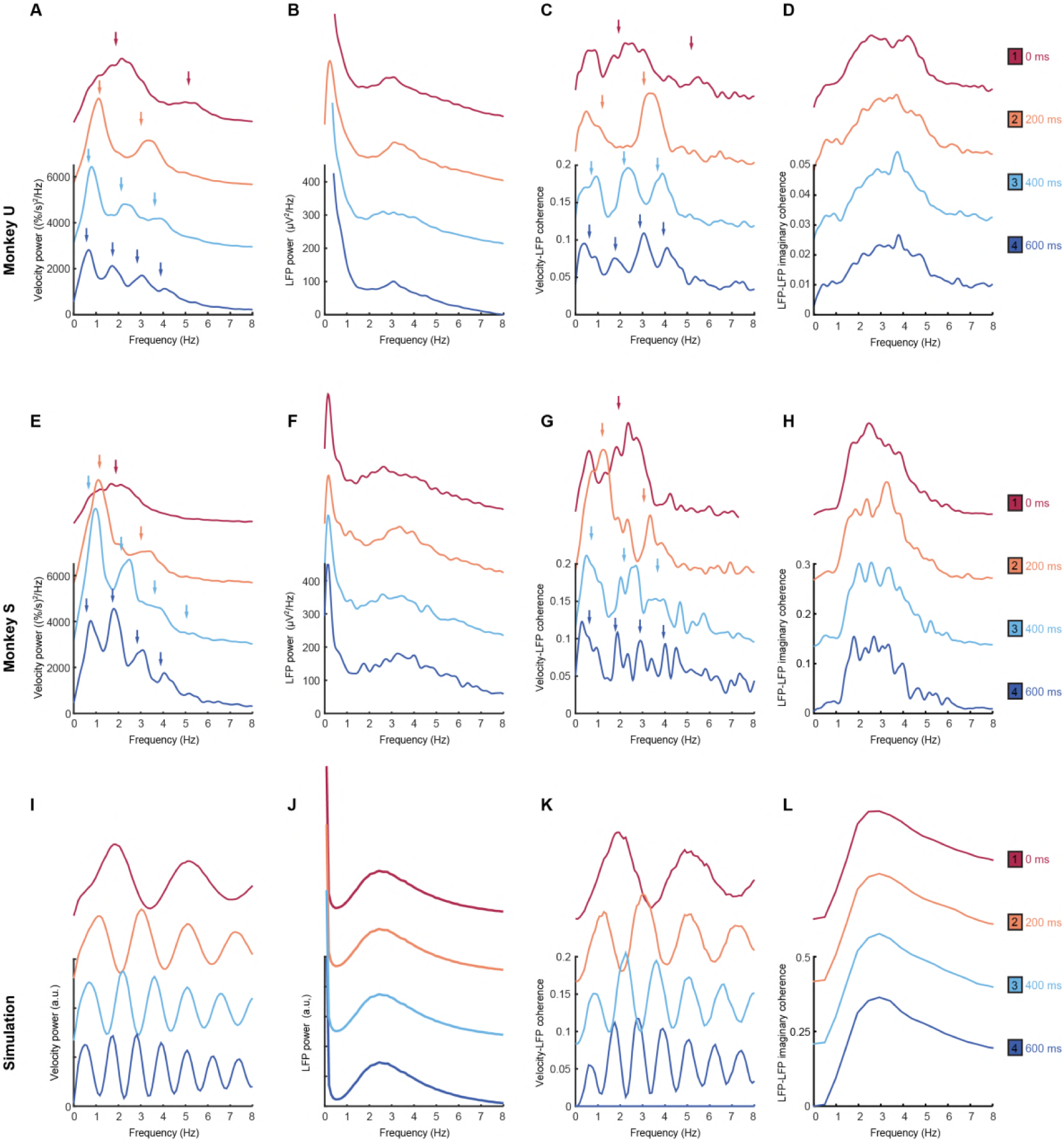
Frequency-domain analysis reveals delay-dependent and delay-independent spectral features. (A) Power spectrum of radial cursor speed with 0–600 ms feedback delay. Traces have been off-set for clarity. Arrows indicate expected frequencies of peaks from OFC model. Data from Monkey U. (B) Average power spectrum of M1 LFPs. (C) Average coherence spectrum between radial cursor speed and all M1 LFPs. (D) Average imaginary coherence spectrum between all pairs of M1 LFPs. (E-H) As above, but for Monkey S. (I-L) Simulated power and coherence spectra produced by the OFC model.

### Feedback delays dissociate intrinsic and extrinsic contributions to intermittency

With increasing feedback delays, submovement peaks in monkeys (Fig. 6A,E) exhibited a pattern similar to that seen with human subjects. The fundamental frequency was reduced, while odd harmonics grew more pronounced as they came below about 4 Hz. Moreover, coherence spectra between cursor velocity and LFP (Fig. 6C,G) revealed peaks at both fundamental and harmonic frequencies. Surprisingly however, the power spectrum of the LFP (Fig. 6B,F) was unaffected by feedback delay, with a single broad peak in the delta band persisting throughout. Moreover, imaginary coherence spectra between pairs of LFPs were also unchanged (Fig. 6D,H). These results appear incompatible with the hypothesis that motor cortical oscillations drive movement intermittency, and instead demonstrate a dissociation between delay-dependent submovements and the conserved rhythmicity of LFPs.

We next identified submovements from peaks in the radial cursor speed, in order to examine the temporal profile of their associated LFPs. Submovement-triggered averages (SmTAs) of LFPs exhibited multiphasic potentials around the time of movement, as well as a second feature following submovements with a latency that depended on extrinsic delay (Fig. 7A, Fig. S6A). This feature was revealed more clearly by reducing the dimensionality of the LFPs with PCA (Fig. 7B, Fig. S6B). Note that if submovements reflect interference between stochastic motor errors and feedback corrections, a submovement in the positive direction can arise from two underlying causes. First, it may be a positive correction to a preceding negative error. In this case, cortical activity associated with the feedback correction should occur around time zero. Second, the submovement may itself be a positive error which is followed by a negative correction, and the associated cortical activity will hence be delayed by the feedback latency. Since the SmTA pools submovements arising from both causes, this accounts for two features with opposite polarity separated by the feedback delay. Note also that SmTAs of cursor velocity similarly overlay (negative) tracking errors preceding (positive) feedback corrections, and (negative) feedback corrections following (positive) tracking errors, evident as symmetrical troughs on either side of the central submovement peak (Fig. 7C, Fig. S6C).

**Figure 7.**
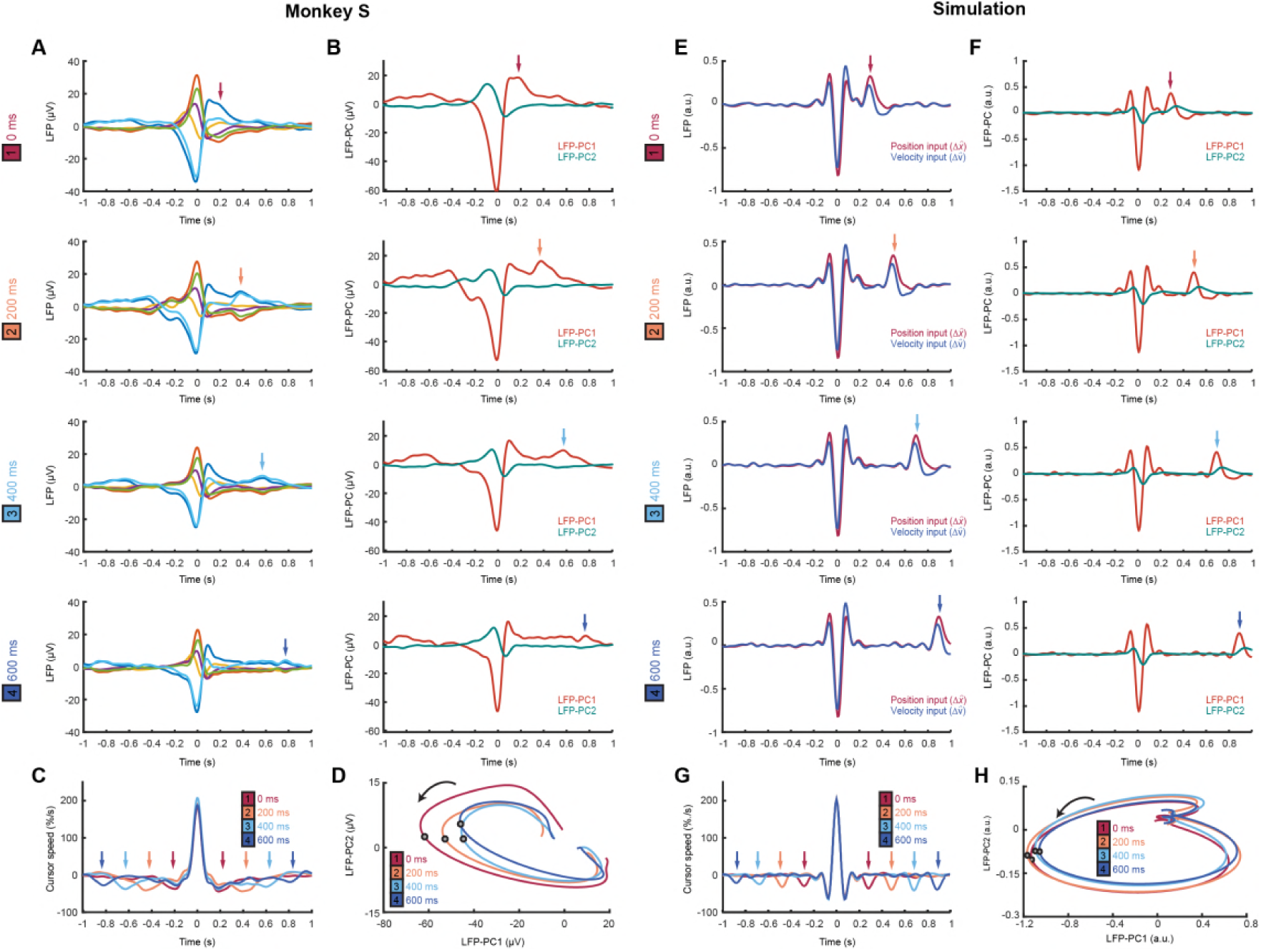
Submovement-triggered averages of M1 LFPs. (A) Average low-pass filtered LFPs from M1, aligned to the peak speed of submovements with 0–600 ms feedback delay. Arrows indicate second feature following submovement by an extrinsic delay-dependent latency. Data from Monkey S. See also Figure S6. (B) Average of first two LFP-PCs aligned to submovements. (C) Average low-pass filtered cursor speed, aligned to submovements. Arrows indicate symmetrical velocity troughs at extrinsic delay-dependent latencies. (D) Average submovement-triggered LFP-PC trajectories, plotted over 200 ms either side of the time of peak submovement speed (indicated by circles). (E-H) Simulated submovement-triggered averages produced by the OFC model.

Importantly however, LFP oscillations around the time of submovements appeared largely unaffected by delay. To visualize this, we projected the SmTAs of multichannel LFPs onto the same PC plane. For all delay conditions, LFPs traced a single cycle with the same directional of rotation and comparable angular velocity (Fig. 7D, Fig. S6D). The period of these cycles (approx. 300 ms) matched the frequency of imaginary coherence between LFPs (approx. 3 Hz), as expected since signals with a consistent phase difference will be orthogonalized by PCA and appear as cyclical trajectories in the PC plane. In other words, although the precise frequency of submovements depends on extrinsic delays in visual feedback, the constant frequency of associated LFP cycles reveals conserved intrinsic dynamics within population activity in the motor cortex. Note also that the resonant frequency of these dynamics matches the intrinsic filtering of feedback responses observed in our human experiments.

### Modelling submovement-related LFP cycles and delta oscillations in sleep

These various observations can be understood using the same computational model that explained our human behavioral data (Figure S7). For simplicity, we simulated two out-of-phase components within the LFP from the total synaptic input to each of the two neural population in the state estimator. We also added common low-frequency background noise to represent volume conduction from distant sources. The simulated LFPs exhibited a broad (delay-independent) spectral peak arising from the dynamics of the recurrent network (Fig. 6J). By contrast, the resultant cursor velocity comprised the summation of motor noise and (delayed) feedback corrections, and therefore contained sharper (delay-dependent) spectral peaks due to constructive/destructive interference (Fig. 6I). Note however that coherence was nonetheless observed between LFPs and cursor velocity (Fig. 6K). Time-domain SmTAs of the simulated data also reproduced features of the experimental recordings, including delay-dependent peaks/troughs reflecting extrinsic feedback delays (Fig. 7E-G). Meanwhile, the conserved intrinsic dynamics coupling simulated neural populations resulted in consistent cyclical LFP trajectories around the time of movement (Fig. 7H) and an imaginary cross-spectrum with a single delay-independent resonance (Fig. 6L).

Finally we examined whether the model could also account for cortical oscillations in the absence of behavior. Previously we have described a common dynamical structure within both cortical cycles during movement and low-frequency oscillations during sleep and sedation (Hall *et al.,* 2014). In particular, K-complex events under ketamine sedation (Fig. 8A), thought to reflect transitions between down- and up-states of the cortex, are associated with brief bursts of delta oscillation (Fig. 8B) (Amzica and Steriade, 1997). The relative phases of multichannel LFPs aligned to these events matches those seen during submovements (Fig. 8D,E). As a result, when projected onto the PC plane, LFPs trace similar cycles during both K-complexes (Fig. 8C) and submovements (Fig. 8F). We modelled the sedated condition by disconnecting motor and sensory connections between the feedback controller and the external world, instead providing a pulsatile input to the state estimator simulating a down-to up-state transition (Fig. 8G). Effectively, transient excitation of the state estimator elicited an impulse response reflecting its intrinsic dynamics. The simulated LFPs generated a burst of delta-frequency oscillation around the K-complex (Fig. 8H) which resembled submovement-related activity (Fig. 8J,K). Projecting this activity onto the same PC plane revealed consistent cycles during simulated K-complexes (Fig. 8I) and submovements (Fig. 8L). Thus it appears that our computational model incorporating the intrinsic dynamics of motor cortical networks could also account for the conserved structure of low-frequency LFPs during movement and delta oscillations in sleep.

**Figure 8.**
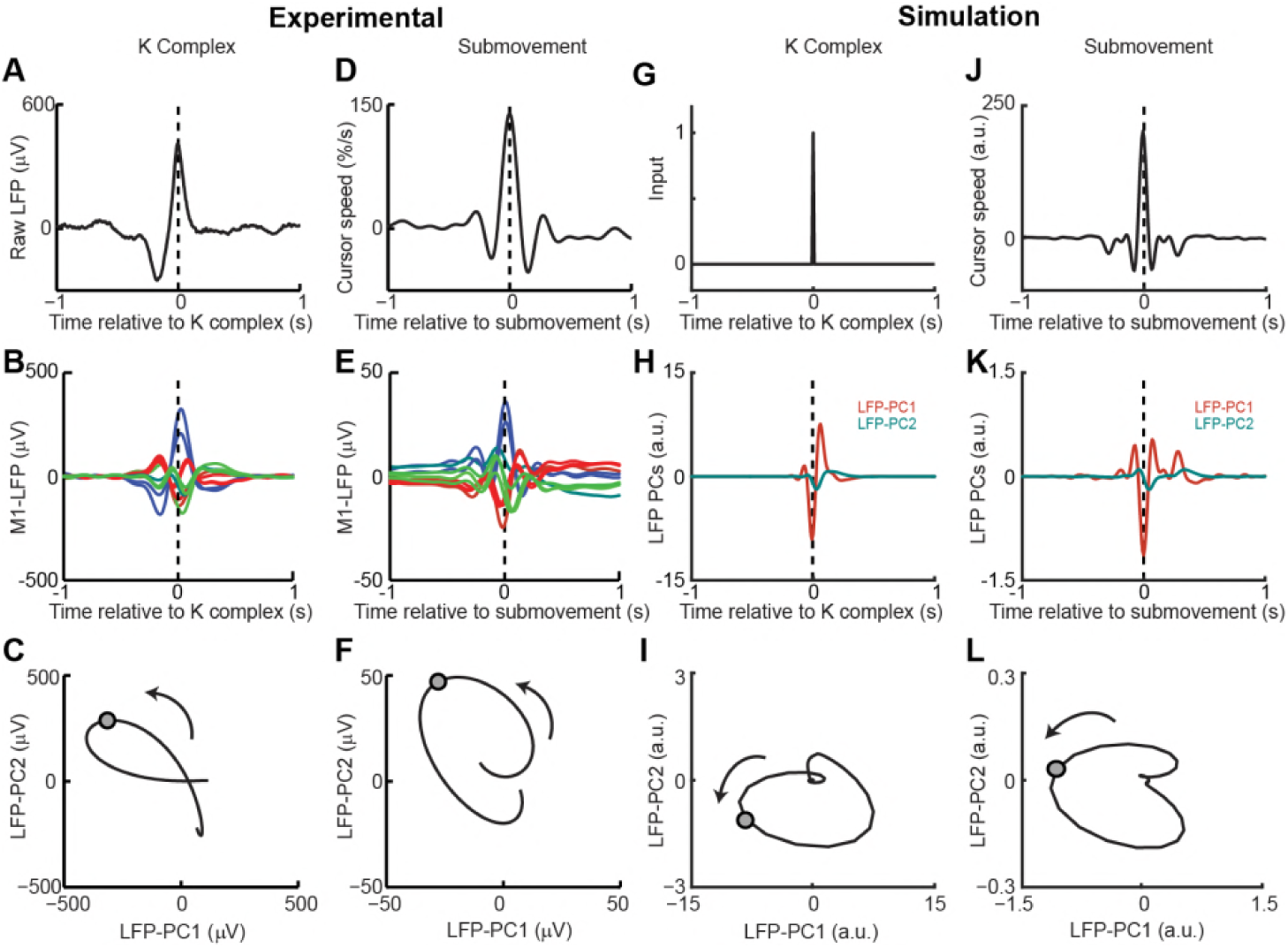
Simulated LFP dynamics during movement and sedation. (A) K-complex events in LFP from M1 recorded under ketamine sedation. (B) Average low-pass filtered multichannel LFPs aligned to K-complex events. LFPs are color-coded according to phase relative to submovements, but exhibit a similar pattern relative to K complexes. (C) Average LFP-PC trajectories aligned to K-complexes, plotted over 200 ms either side of the time of K-complex (indicated by circle), using the PC plane calculated from recordings during awake behavior. (D) Average cursor speed aligned to the peak speed of submovements. (E) Average low-pass filtered multichannel LFPs aligned to submovements. (F) Average submovement-triggered LFP-PC trajectories, plotted over 200 ms either side of the time of submovements (indicated by circle). (G) A K-complex under sedation is simulated by an impulse excitation of the OFC model, without connection to the external world. (H) Impulse response of the simulated LFP-PCs. (I) LFP-PC trajectories associated with simulated K-complexes. (J) Simulated submovement-triggered average cursor speed from the OFC model with no feedback delay. (K) Simulated submovement-triggered average LFP-PCs. (L) Simulated submovement-triggered LFP-PC trajectories. Panels A-F reproduced from Figure 4A,C,D in Hall et al. (2014) under the CC BY 3.0 license (https://creativecommons.org/licenses/by/3.0/).

## Discussion

### Submovement kinematics are influenced by both extrinsic and intrinsic dynamics

Previous theories of intermittency have focused on either extrinsic or intrinsic explanations for the regularity of submovements, but little consensus has emerged over this fundamental feature of movement. There is good evidence for a common low-frequency oscillatory structure to motor cortex activity across multiple behavioral states (Churchland *et al.,* 2012; Hall *et al.,* 2014; Russo *et al.,* 2018) but also an influence of feedback delays on submovement timing (Miall, 1996). Experimentally manipulating visual feedback with artificial time delays and spatial perturbations allowed us to dissociate both contributions to submovement kinematics. We found that precise frequencies of submovement peaks were determined by constructive and destructive interference between broad-band motor errors and continuous, delayed feedback corrections. However, these peaks were embedded within a delay-independent envelope that arose from intrinsic filtering of feedback corrections. The dissociation of extrinsic and intrinsic dynamics was also evident in cortical LFPs during tracking movements. Both delay-dependent feedback corrections and delay-independent cycles were observed in submovement-triggered averages of LFPs. Moreover, while coherence between LFPs and cursor movement exhibited delay-dependent spectral peaks, the imaginary coherence between multichannel LFPs revealed a consistent dynamical structure across behaviors.

These apparently contradictory results could be explained by an OFC model that implemented state estimation via a steady-state Kalman filter to separate process (motor) noise from measurement (sensory) noise. One free parameter was tuned to achieve correspondence between simulated and experimental data, namely the ratio of process to measurement noise which determined the intrinsic resonance frequency around 2-3 Hz. It would be interesting in future to vary these noise characteristics experimentally (e.g. by artificially degrading visual acuity or by extensively training subjects) and examine the effect on perturbation responses. One possible outcome would be a change to the observed resonance, although this seems to contradict the ubiquity of 2-3 Hz cortical dynamics. Alternatively there may be other computational advantages to maintaining a consistent cortical rhythm. For example, it is notable that 2-3 Hz intrinsic dynamics matched the frequency of the primary submovement peak under unperturbed external feedback conditions, thus accentuating the fundamental submovement frequency around 2 Hz while suppressing higher harmonics. This may be beneficial in allowing other aspects of the visuomotor machinery to be synchronized to a single rhythm, for example eye movements which are influenced by hand movement during tracking tasks (Koken and Erkelens, 1992).

### Modelling isometric visuomotor tracking

Several further assumptions of our modelling warrant discussion. First, to prevent control instabilities associated with feedback delays we incorporated an accurate forward model of the (delayed) sensory consequences of motor commands within a Smith Predictor architecture. We did not include adaptive processes to calibrate the delay model, but this could be readily achieved by minimizing discrepancies between an efference copy of motor commands and observed cursor movements. The accuracy of tracking performance under different delay conditions (without cursor perturbations) suggests that subjects could readily adapt such a forward model and this role has previously been ascribed to the cerebellum (Miall *et al.,* 1993b; Streng *et al.,* 2018). By contrast, to account for the delay-independent perturbation responses, we maintained the same intrinsic cortical dynamics throughout, even though an optimal state estimator should similarly incorporate knowledge of feedback delays (see Equ. 5). Note however that adaptation of the state estimator presents a harder computational problem, since no available signals directly relate the state of the external world to delayed sensory information. Even when we imposed predicTable Sinusoidal perturbations, we saw no evidence that subjects learnt to compensate for feedback delays by altering the timing of their corrective responses within a single trial. Nevertheless, it would again be interesting to examine whether state estimator dynamics might adapt on a slower time-scale after extensive training with delayed feedback.

Finally, we were puzzled that force amplitude responses to cursor perturbations were uniformly less than unity, which initially appears suboptimal for rejecting even slow perturbations. We first considered that proprioceptive information (which is in conflict with vision during cursor perturbations) might cause subjects to underestimate the true displacement of the cursor. However, sub-unity amplitude responses were also observed in separate experiments (not shown) when sinusoidal displacements were added to the target position. In this situation there was no discrepancy between vision and proprioception, yet subjects consistently undershot corrections to all but the lowest frequency perturbations (even in the absence of any delay). In our OFC model we instead reduced amplitude responses by penalizing large changes to the motor command. This cost function was minimized by proportional-integral (PI) control, which has been used in the past to model human movement (Kleinman, 1974). It is more common in current optimal control models to apply cost functions that penalize the absolute motor command leading to proportional feedback policies (Todorov and Jordan, 2002), under the assumption that this minimizes signal-dependent noise in muscles (Jones *et al.,* 2002). However, the trajectory variability observed in our isometric tracking task appeared more correlated with large changes in finger forces rather than force magnitude (Figure S8), providing empirical support for our choice of cost function. Derivative-dependent motor noise was also evident as increased variability at high frequencies in our feedforward task (Figure S5). Since submovements result from constructive interference between tracking errors and feedback corrections, derivative-dependent motor noise also provides a counterintuitive but necessary explanation for why the amplitude of submovements increases with target speed (Figure S1). Increased intermittency cannot be a direct consequence of faster target motion, since the frequency content of this motion is nevertheless low by comparison to submovements. Rather, faster tracking requires a larger change in the motor command, leading to increased broad-band motor noise which, after constructive interference with feedback corrections, results in more pronounced peaks at submovement frequencies.

### State estimation by motor cortical population dynamics

PCA of multichannel LFPs in monkey motor cortex revealed two underlying components, which we interpret as arising from distinct but coupled neural populations. The cyclical movement-related dynamics of these components resembled those described for M1 firing rates (Churchland *et al.,* 2012), which have previously been implicated in feedforward generation of movement. Specifically, it was proposed that preparatory activity first develops along ‘output-null’ dimensions of the neural state space before, at movement onset, evolving via intrinsic dynamics into orthogonal ‘output-potent’ dimensions that drive muscles (Churchland *et al.,* 2010). However, this purely feedforward view cannot account for our isometric tracking data, since manipulation of feedback delays dissociated delay-dependent submovements from delay-independent rotational dynamics. Instead we interpret these intrinsic dynamics as implementing a state estimator during continuous feedback control. We used Newtonian dynamics to construct a simple two-dimensional state transition model based on both the cursor-target discrepancy and its first derivative. While this undoubtedly neglects the true complexity of muscle and limb biomechanics, simulations based on this plausible first approximation reproduced both the amplitude response and phase delay to sinusoidal cursor perturbations in humans, and the population dynamics of LFP cycles in the monkey. Note that this account also offers a natural explanation of why preparatory and movement-related activity lies along distinct state-space dimensions, since the static discrepancy present during preparation is encoded differently to the changing discrepancy that exists during movement. At the same time, the lawful relationship between discrepancy and its derivative couples these dimensions within the state estimator and is evident as consistent rotational dynamics across different tasks and behavioral states.

It may seem unusual to ascribe the role of state estimation to M1 when this function is usually attributed to parietal (Mulliken *et al.,* 2008) and premotor areas (where rotational dynamics have also been reported, albeit at a lower frequency (Churchland *et al.,* 2012; Hall *et al.,* 2014). We suggest that the computations involved in optimal visuomotor tracking are likely distributed across multiple cortical areas including (but not limited to) M1, with local circuitry reflecting multiple dynamical models of the various sensory and efference copy signals that must be integrated for accurate control. Indeed, while we neglected to model the computations involved in accurately estimating our slow and predictable target motion, state estimation using Kalman filters has also been suggested as a mechanism by which the visual system can estimate the position of moving visual stimuli (Kwon *et al.,* 2015).

An alternative explanation for consistent rotational dynamics has recently been proposed by Russo *et al.* (2018), based on the behavior of recurrent neural networks trained to produce different feed-forward muscle patterns whilst minimizing ‘tangling’ between neural trajectories. It is interesting to compare this with our OFC-based interpretation, since both are motivated by the problem of maintaining accurate behavior in the presence of noise. Minimizing tangling leads to network architectures that are robust to intrinsic noise in individual neurons, while OFC focusses on optimizing movements in the face of unreliable motor commands and noisy sensory signals. Given this conceptual link, it is perhaps unsurprising if recurrent neural network approaches learn implementations of computational architectures such as Kalman filters that minimize the influence of noise on behavior. In future it may be productive to incorporate sensory feedback into recurrent neural network models of movement, as well as including intrinsic sources of neural noise in optimal control models. The convergence of these frameworks may further help to reveal how computational principles are implemented in the human motor system.

## Acknowledgements

We thank Jenifer Tulip and Norman Charlton for technical assistance. This work was supported by the Indonesia Endowment Fund for Education (S-2648/LPDP.3/2014), the Medical Research Council (K501396) and the Wellcome Trust (106149).

## Author contributions

DS, KA and AJ designed the study. DS collected the human data. WX and TMH collected the monkey data. DS, FG and AJ developed the computational model. DS and AJ wrote the manuscript with contributions from all other authors.

## Declaration of competing interests

The authors declare no competing interests.

## Materials and Methods

### Human experiments

#### Subjects

Based on pilot studies we decided in advance to use a sample size of eight subjects in each experiment. In total, we recruited 11 adult subjects in total at the Institute of Neuroscience, Newcastle University. Eight subjects (3 females; age 23–33; 1 left-handed) participated in both Experiment 1 (feedback delay) and Experiment 2 (feedback delay and spatial perturbation). Eight subjects (3 females; age 23– 33; all right-handed) participated in Experiment 3 (feedforward task); 6 of these subjects also participated in experiments 1 and 2. Eight subjects (3 females; age 23–33; all right-handed) participated in the experiment shown in Figure S8; 7 out of these subjects also participated in Experiment 3. All experiments were approved by the local ethics committee at Newcastle University and performed after informed consent, which was given in accordance with the Declaration of Helsinki.

#### Human tracking task

Subjects tracked a (red) target on a computer monitor by exerting bimanual isometric index finger forces on two sensors (FSG15N1A; Honeywell). The target underwent uniform, slow circular motion with a pseudorandom order of clockwise and anticlockwise directions across trials. Finger forces were sampled at 50 samples/s (USB-6343; National Instruments) and mapped to (yellow) cursor position by projecting onto two diagonal screen axis. In addition, a feedback delay (*τ*_ext_) was interposed between force and cursor movement. The feedback delay was kept constant through the duration of each trial (lasting 20 s). We express screen coordinates in terms of the radius of target motion, *r*_target_ = 100%. Tracking the target rotation thus required generating sinusoidal motion in the range of -100% to +100%, corresponding to finger forces of 0 to 3.26N, with a 90° phase-shift between each hand. At the end of each trial subjects were given a numerical score from 0-1000 indicating how accurately they tracked the target. Subjects were instructed to attempt to maximize this score, which was calculated as:

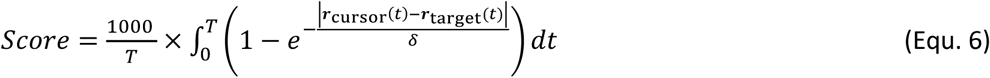

where ***r***_cursor_(*t*) and ***r***_target_(*t*) are the 2D positions of the cursor and target respectively, and *δ* = 50%. Apart from the experiment shown in Figure S8, all experiments used a frequency of target rotation, *f*_target_ = 0.2 rotations per second.

Experiment 1 used five delay conditions (*τ*_ext_ = 0, 100, 200, 300, or 400 ms). Subjects performed a total of 70 trials, comprising 14 of each condition presented in pseudorandom order.

For Experiment 2, spatial perturbations were added to the cursor position as well as time delays. The perturbations were equivalent to sinusoidal modulation of the target angular velocity, but were instead added to the cursor. Expressed in polar coordinates ***r*** = ⟨*r,* ∠*θ*⟩ relative to the center of the screen, the target and cursor positions were thus given by:

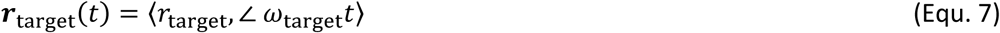

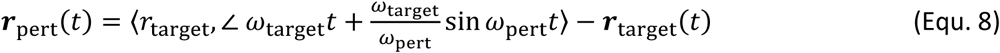

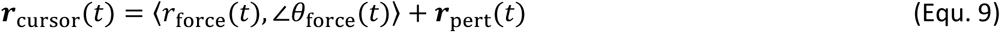

where *ω*_target_ = 2*πf*_target_ is the angular velocity of the target around the centre of the screen, *ω*_pert_ = 2*πf*_pert_ is the angular frequency of the perturbation, and ⟨*r*_force_(*t*), ∠*θ*_force_(*t*)⟩ is the unperturbed cursor position calculated from the subject’s forces at time *t* - *τ*_ext_.

Kinematic analyses were based on the time-varying angular velocity of the cursor subtended at the center of the screen:

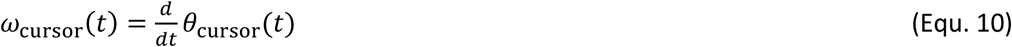

For spatial perturbation experiments, we also calculated the angular velocity of the unperturbed cursor position subtended at the center of the screen:

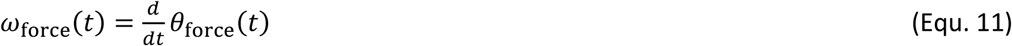

Note that since *r*_force_ ≈ *r*_target_, the perturbation effectively adds a sinusoidal component to the angular velocity of the cursor:

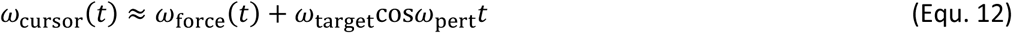

Six different spatial perturbations (*f*_pert_ = 0, 1, 2, 3, 4, 5 Hz) combined with two feedback delays (*τ*_ext_ = 0, 200 ms) yielded 12 conditions. Subjects performed a total of 144 trials, comprising 12 trials per condition in pseudorandom order.

#### Human feedforward task

In Experiment 3, we used a unimanual isometric task in which subjects were asked to make sinusoidal forces with their right index finger. Subjects received visual feedback of the cursor, but no target was shown. Instead subjects were shown two amplitude boundaries to move between, and the frequency of movement was cued with auditory beeps at frequencies of 1, 2, 3, 4 and 5 Hz. Subjects performed a total of 15 trials, comprising three 20 s trials per frequency condition.

## Monkey experiments

### Subjects

We used two purpose-bred female rhesus macaques (monkey S: 6 years old, 6.6 kg; monkey U: 6 years old, 8.8 kg). Animal experiments were approved by the local Animal Welfare Ethical Review Board and performed under appropriate UK Home Office licenses in accordance with the Animals (Scientific Procedures) Act 1986.

### Monkey tracking task

Monkeys moved a 2D computer cursor by generating isometric flexion-extension (vertical) and radial-ulnar (horizontal) torques at the wrist, measured by a 6-axis force/torque transducer (Nano25; ATI Industrial Automation). Centre-out targets were presented at 8 peripheral positions in a pseudorandom order. Targets were positioned at 70% of the distance to the screen edge (100% corresponding to torque of 0.67 Nm). The diameter of the target and cursor ranged between 14-36%. A successful trial required maintaining an overlap between cursor and peripheral target for 0.6 s after which the monkeys returned to the center of the screen to receive a food reward. Visual feedback of the cursor was delayed by *τ*_ext_ = 0, 200, 400, 600 ms throughout separate blocks of 50-70 trials each. Monkey S performed the task with the right hand. Monkey U initially used the right hand and was then retrained for a second period of data collection with the left hand.

### LFP recording

LFPs were recorded using custom arrays of 12 moveable 50 μm diameter tungsten microwires (impedance ^∼^200 kΩ at 1 kHz) chronically implanted in contralateral wrist area of M1 under sevoflurane anesthesia with postoperative analgesics and antibiotics. Head-free recordings were made using unity-gain headstages followed by wide-band amplification and sampling at 24.4 kilosamples/s (System 3; Tucker-Davis Technologies). LFPs were digitally low-pass filtered at 200 Hz and recorded at 488 samples/second.

Analysis of kinematics and neural data was performed on data recorded over 8 different days comprising of 56 task blocks in Monkey S (no delay: 24 blocks; 200 ms delay: 13; 400 ms delay: 13; 600 ms delay: 6), and 89 recording days comprising of 356 task blocks in Monkey U (no delay: 89; 200 ms delay: 89; 400 ms delay: 89; 600 ms delay: 89). Each task block comprised 50 (monkey S) or 70 trials (monkey U).

## Analysis Methods

### Human data analysis

Spectral analysis used fast Fourier transforms (FFTs) performed on non-overlapping 512 sample-point windows (approx. 10s) taken from the middle of each trial. Submovement peaks in the power spectra were measured after smoothing with a seven-point moving-average.

For perturbation experiments, we additionally defined two complex transfer functions *H*_cursor_ and *H*_force_:

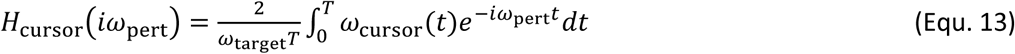

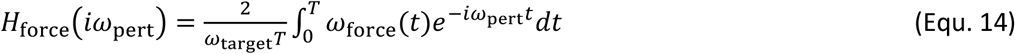

Cursor and force amplitude responses to perturbations were calculated as the magnitude of the corresponding transfer functions, and the intrinsic phase delay of force responses was given by:

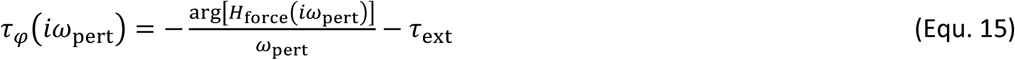

Additionally, tracking performance was quantified off-line using the root-mean-squared Euclidean distance between cursor and target.

### Monkey data analysis

We differentiated the magnitude of the absolute 2D torque (expressed as a percentage of the distance to the edge of the screen) to obtain the radial cursor velocity. LFP channels were subjected to visual inspection to reject noisy channels prior to mean-subtraction. For time-domain analysis, LFPs and cursor velocities were low-pass filtered at 10 Hz. Submovements were defined as a peak radial cursor speed exceeding 100%/s (monkey S) and 150%/s (monkey U). For frequency-domain analysis, we took unfiltered sections of 1024 sample points from each trial (approx. 1.5 s before to 0.5 s after the end of the peripheral hold period). We subtracted the trial-averaged profile from each section before concatenating to yield long data sections without any consistent low-frequency components related to the periodicity of the task. FFTs were calculated with overlapping Hanning windows (2^14^ sample points ≈ 34 s; 75% overlap), from which we derived the following spectra:

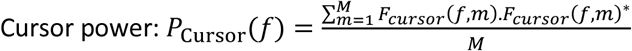

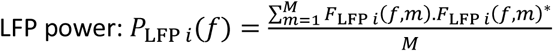

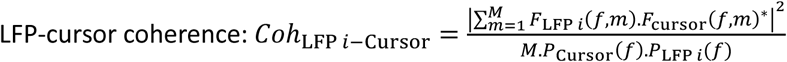

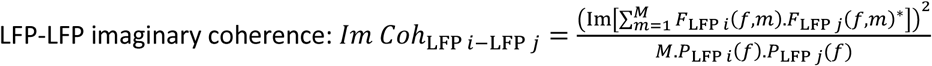

where *F*_LFP i_(*f, m*) and *F*_Cursor_(*f, m*) represent Fourier coefficients at frequency *f* and window *m* = (1.\*M*) from LFP channel *i* and cursor velocity respectively. All spectra were smoothed with a 16-point Hanning window. In addition, LFP power and LFP-cursor coherence were averaged across all LFP channels, while LFP-LFP imaginary coherence was averaged over all pairs of LFPs.

## Modelling

Although both human and monkey tasks involved 2D isometric control, for simplicity we modelled only a 1D controller and assumed a one-to-one mapping from control signal, u_*k*_ to position, *x*_k_. We neglected target motion and designed the controller to minimize the influence of stochastic motor errors using delayed, noisy feedback of position. We set the model time step Δ*t* = 0.01 s, intrinsic feedback delay *τ*_int_ = 0.26 s, and the ratio of process/measurement noise *ρ* = 250 s^-2^ unless otherwise stated. Steady-state Kalman gains were calculated using the function *kalman* in MATLAB, and the resultant discrete time dynamic system (Equ. 4) was implemented by two integrating neuronal populations representing 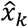 and 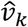, receiving a synaptic input on each time-step equal to:

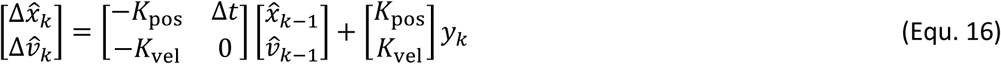

Two LFP components were simulated by normalizing 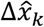 and 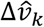 to unity variance, before adding background common noise with a 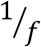 spectrum.

The motor command u_*k*_ was generated on each time step using the Smith Predictor architecture shown in Fig. 3F. Based on our observation that trajectory variability was maximal at times when force output was changing (Figure S8), we used an linear quadratic regulator (LQR) control framework to minimize a quadratic cost function, *J,* incorporating the rate of change in motor command, 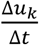:

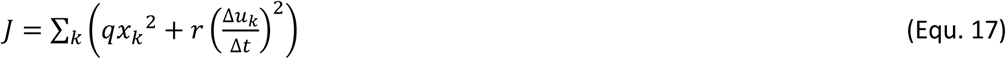

For a state transition matrix in the form:

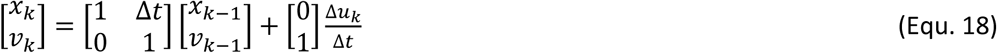

*J* is minimized by a state feedback policy of the form:

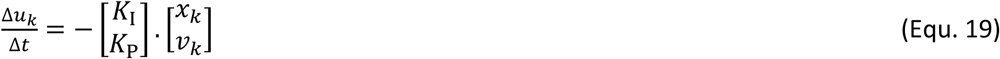

which can be integrated to yield a PI controller:

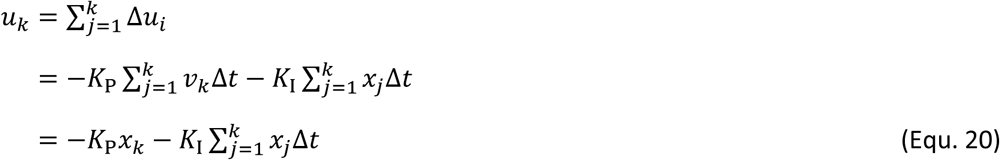

We found the proportional and integral gains *K*_P_ and *K*_I_ using the function *lqr* in MATLAB with *q* = 1 and *r* = Δ*t*^2^. In the full model, this controller acted on the optimal estimate of position, 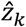, after incorporating the delay feedback loop of the Smith Predictor. Note that the transfer function of a PI controller inside the fast feedback loop of the Smith Predictor is given by (Abe and Yamanaka, 2003):

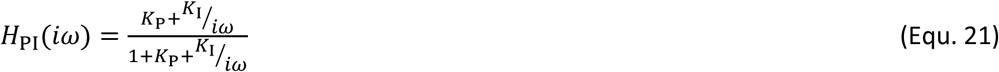

which equals 1 for ω = 0 but tends to 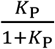 at higher frequencies. Therefore this effectively reduces the response amplitude to perturbations. The full transfer function of the intrinsic dynamics, including time-delay is given by:

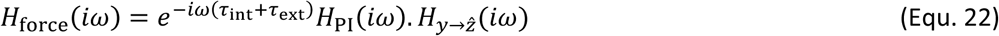

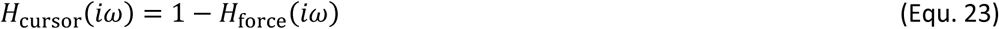

where 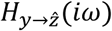 is the transfer function of the Kalman filter relating delayed position measurement to optimal position estimate.

## Data and software availability

Datasets from the human and monkey experiments, sample analysis code and modelling associated with this work are available on Dryad doi:10.5061/dryad.53sq7kn.

## Supplemental Information

**Table S1.**
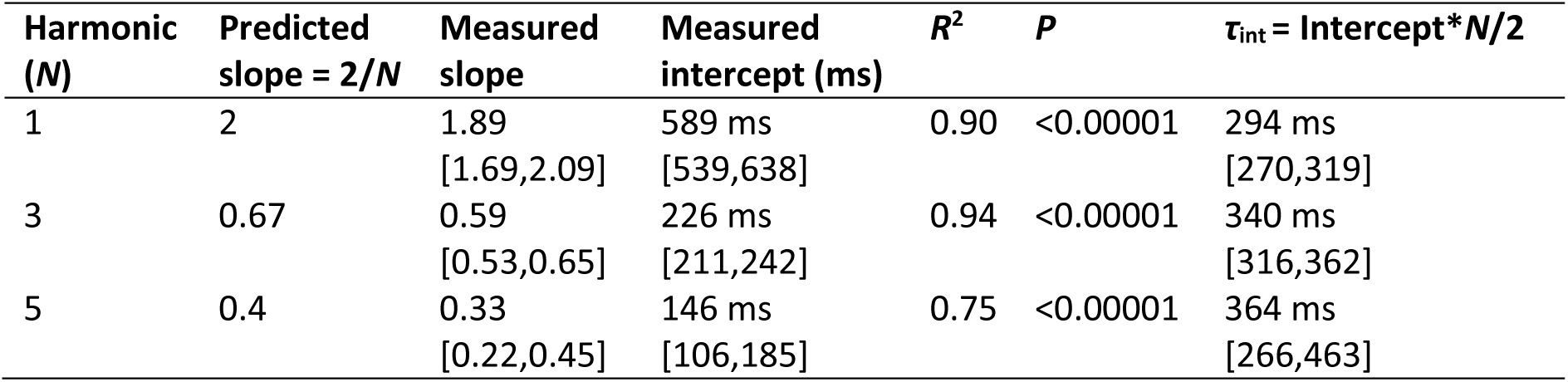
The dependency of submovement period on feedback delay. Shown in the table are the gradients and intercepts of regression lines fitted to each harmonic group in Figure 1E. The time period of each spectral peak was regressed against feedback delay. Shown in square brackets are 95% confidence intervals of these values. Also shown is the estimated intrinsic time delay calculated using Equ. 1.

| Harmonic<br>(*N*) | Predicted<br>slope = $2/N$ | Measured<br>slope | Measured<br>intercept (ms) | $R^2$ | $P$ | $\tau_{\text{int}} = \text{Intercept} * N/2$ |
| --- | --- | --- | --- | --- | --- | --- |
| 1 | 2 | 1.89<br>[1.69,2.09] | 589 ms<br>[539,638] | 0.90 | <0.00001 | 294 ms<br>[270,319] |
| 3 | 0.67 | 0.59<br>[0.53,0.65] | 226 ms<br>[211,242] | 0.94 | <0.00001 | 340 ms<br>[316,362] |
| 5 | 0.4 | 0.33<br>[0.22,0.45] | 146 ms<br>[106,185] | 0.75 | <0.00001 | 364 ms<br>[266,463] |

**Figure S1.**
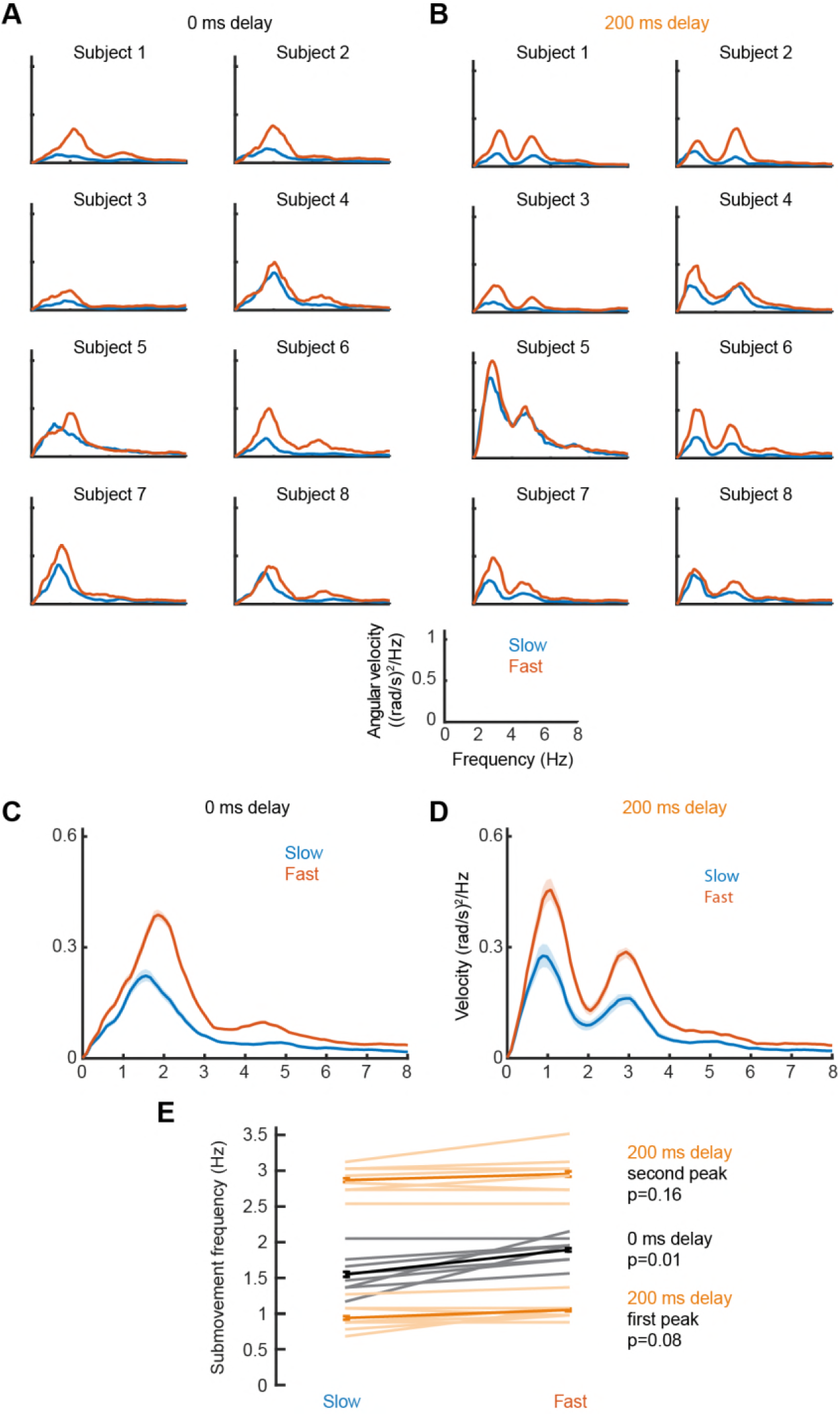
Effect of target speed on movement intermittency. (A) Power spectra of cursor angular velocity for individual subjects with slow (0.1 cycles/s) or fast (0.2 cycles/s) target rotation, and no feedback delay. (B) Power spectra of cursor angular velocity with slow or fast target rotation, and 200 ms feedback delay. (C,D) Average power, showing mean ± s.e.m. for 8 subjects. (E) Average ± s.e.m. frequencies of peak cursor velocity in each condition. P values calculated using a paired t-test.

**Figure S2.**
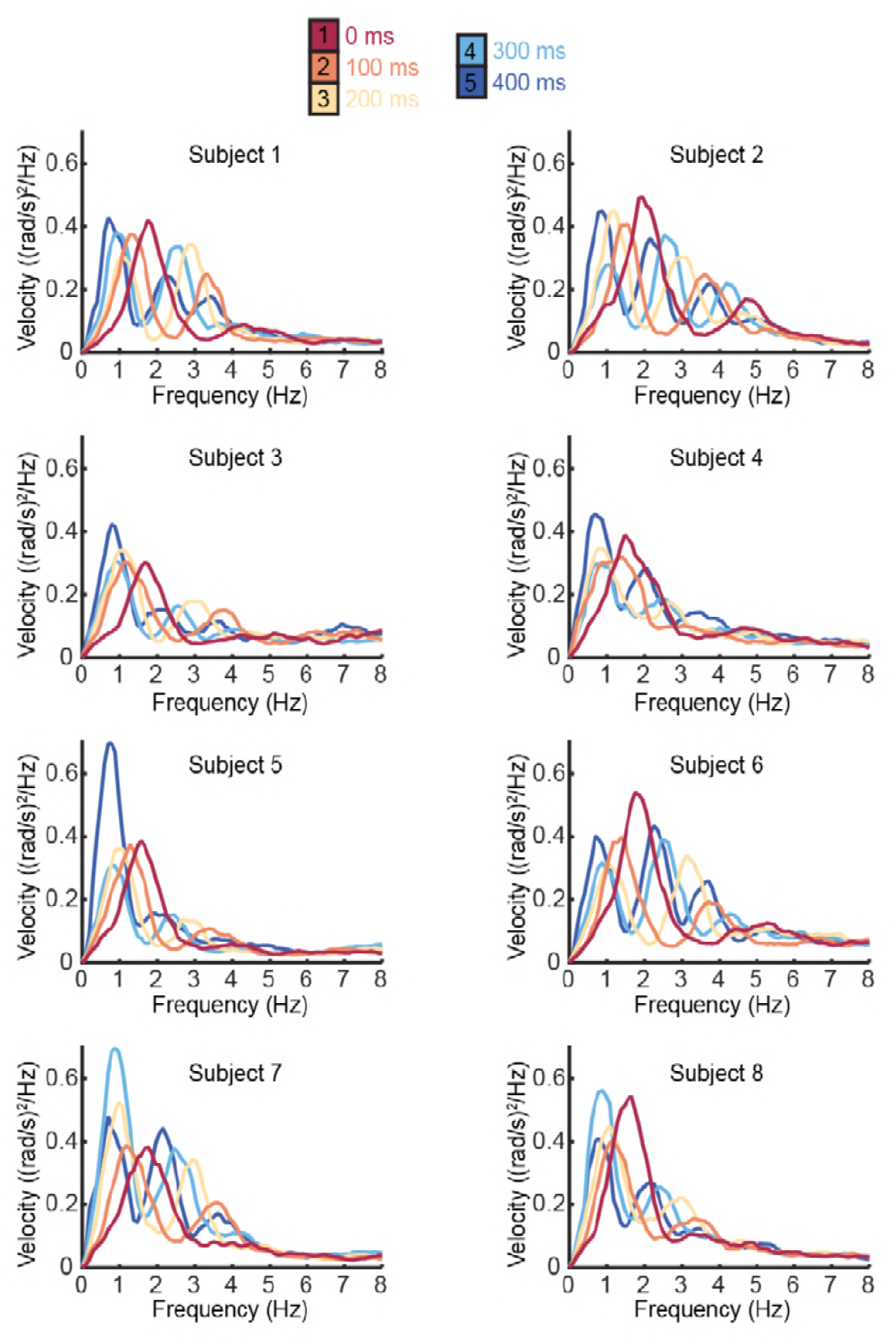
Individual subject power spectra of cursor velocity with different feedback delays. Power spectra of cursor angular velocity for individual subjects with 0–400 ms feedback delay. The average over subjects is shown in Figure 1D.

**Figure S3.**
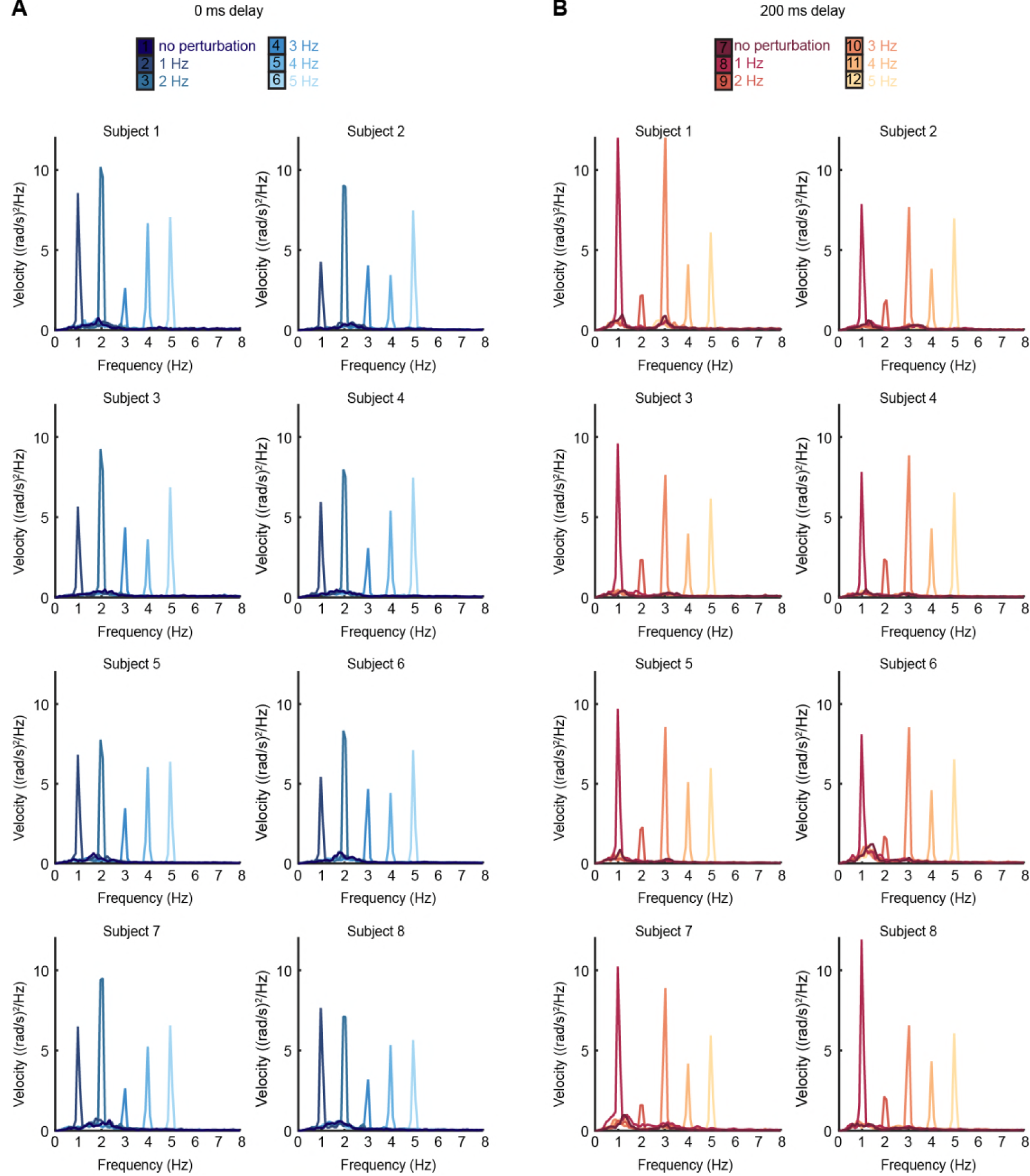
Individual subject power spectra of cursor velocity with perturbations. (A) Power spectra of cursor angular velocity for individual subjects with 1–5 Hz perturbations and no feedback delay. The average over subjects is shown in Figure 2C. (B) Power spectra of cursor angular velocity for individual subjects with 1–5 Hz perturbations and 200 ms feedback delay. The average over subjects is shown in Figure 2D.

**Figure S4.**
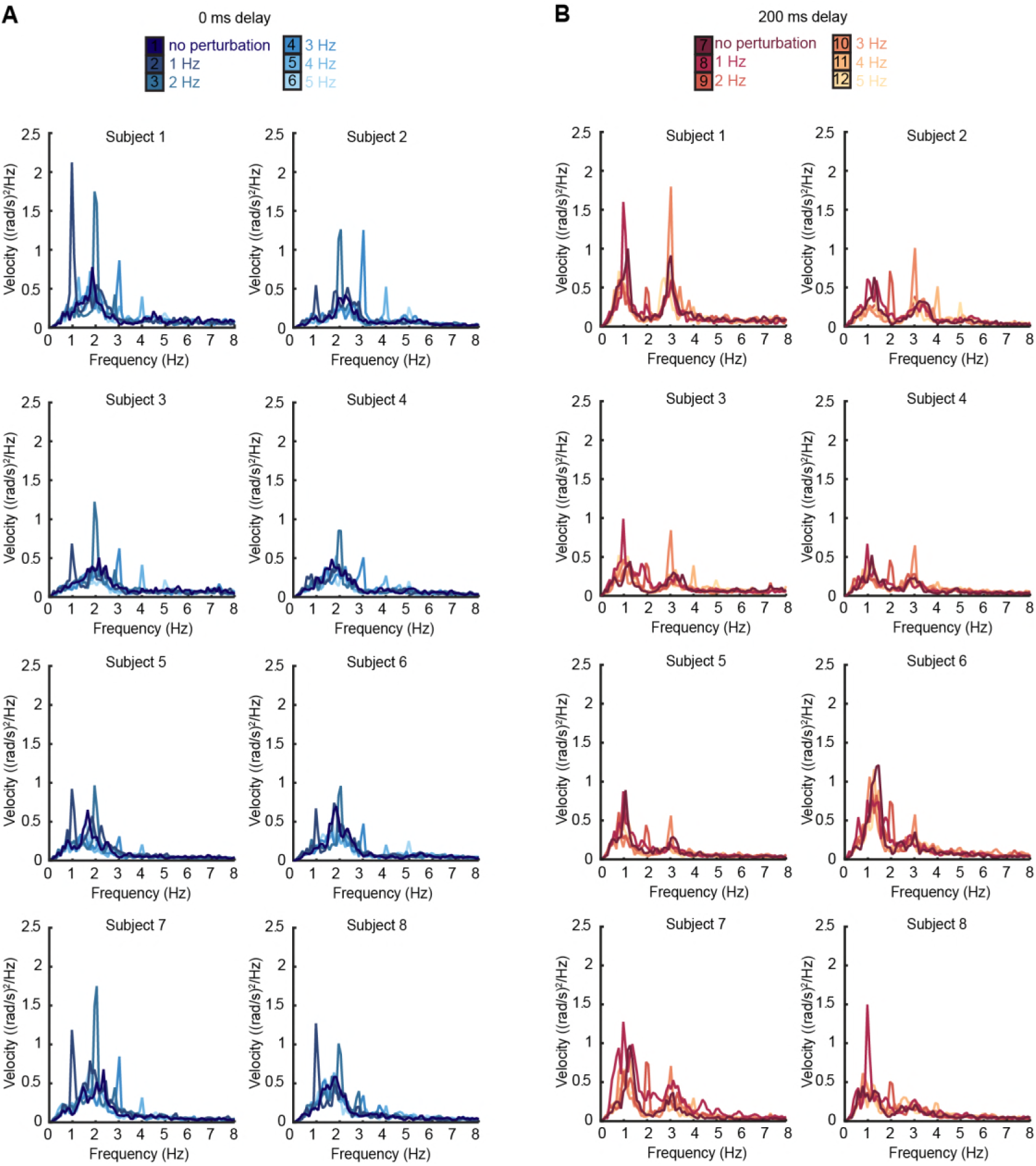
Individual subject power spectra of force velocity with perturbations. (A) Power spectra of force angular velocity for individual subjects with 1–5 Hz perturbations and no feedback delay. The average over subjects is shown in Figure 2F. (B) Power spectra of force angular velocity for individual subjects with 1–5 Hz perturbations and 200 ms feedback delay. The average over subjects is shown in Figure 2G.

**Figure S5.**
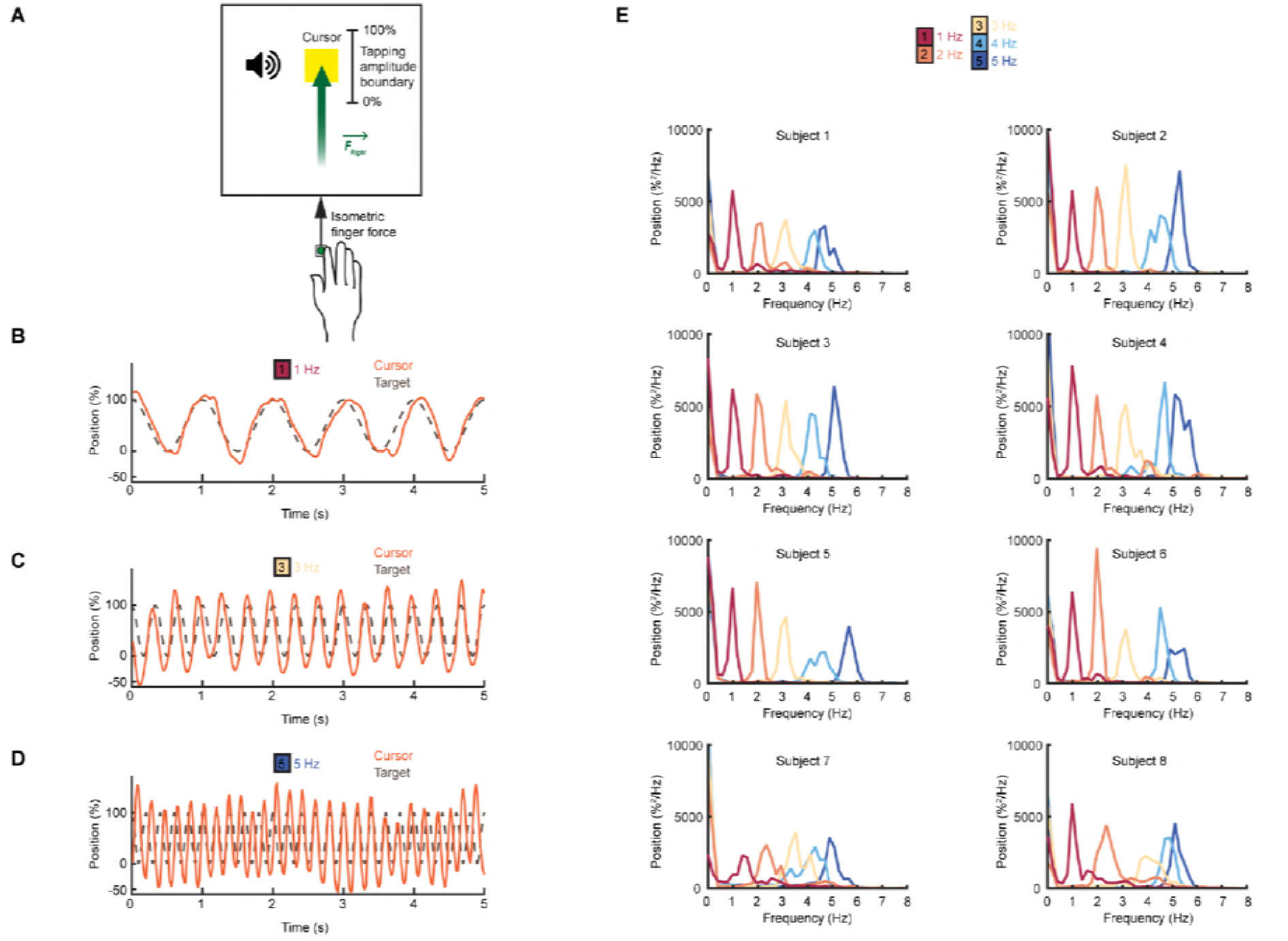
Feedforward task. (A) Schematic of the feedforward isometric task. Subjects generated sinusoidal forces within a set range, at a frequency indicated by an auditory cue. (B-D) Performance of an example subject for frequencies between 1-5 Hz. (E) Power spectrum of force for individual subjects. The average over all subjects is shown in Figure 2J.

**Figure S6.**
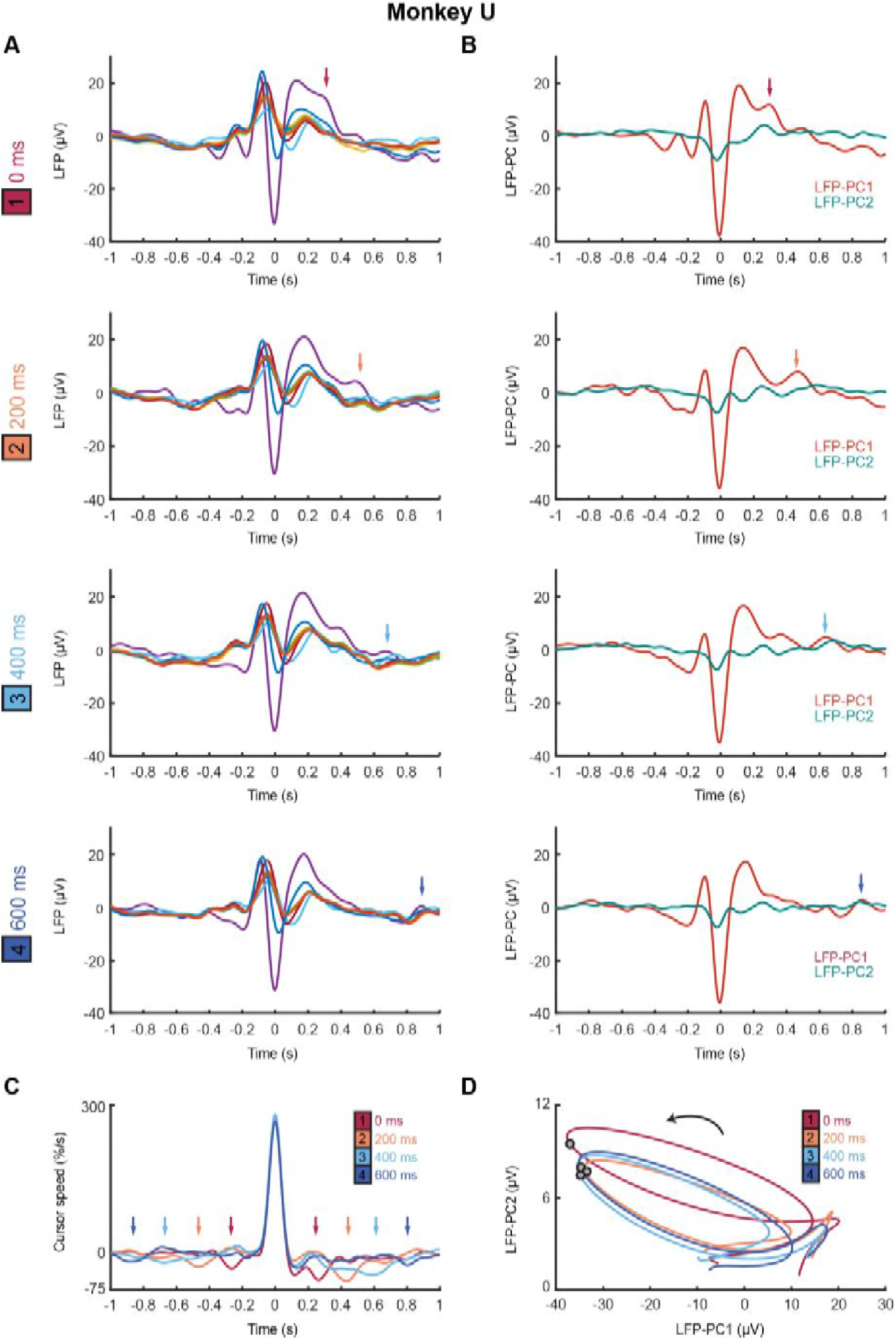
Submovement-triggered averages of M1 LFPs for Monkey U. (A) Average low-pass filtered LFPs from M1, aligned to the peak speed of submovements with 0–600 ms feedback delay. Arrows indicate second feature following submovement by an extrinsic delay-dependent latency. Data from Monkey U. (B) Average of first two LFP-PCs aligned to submovements. (C) Average low-pass filtered cursor speed, aligned to submovements. Arrows indicate symmetrical velocity troughs at extrinsic delay-dependent latencies. (D) Average submovement-triggered LFP-PC trajectories, plotted over 200 ms either side of the time of peak submovement speed (indicated by circles).

**Figure S7.**
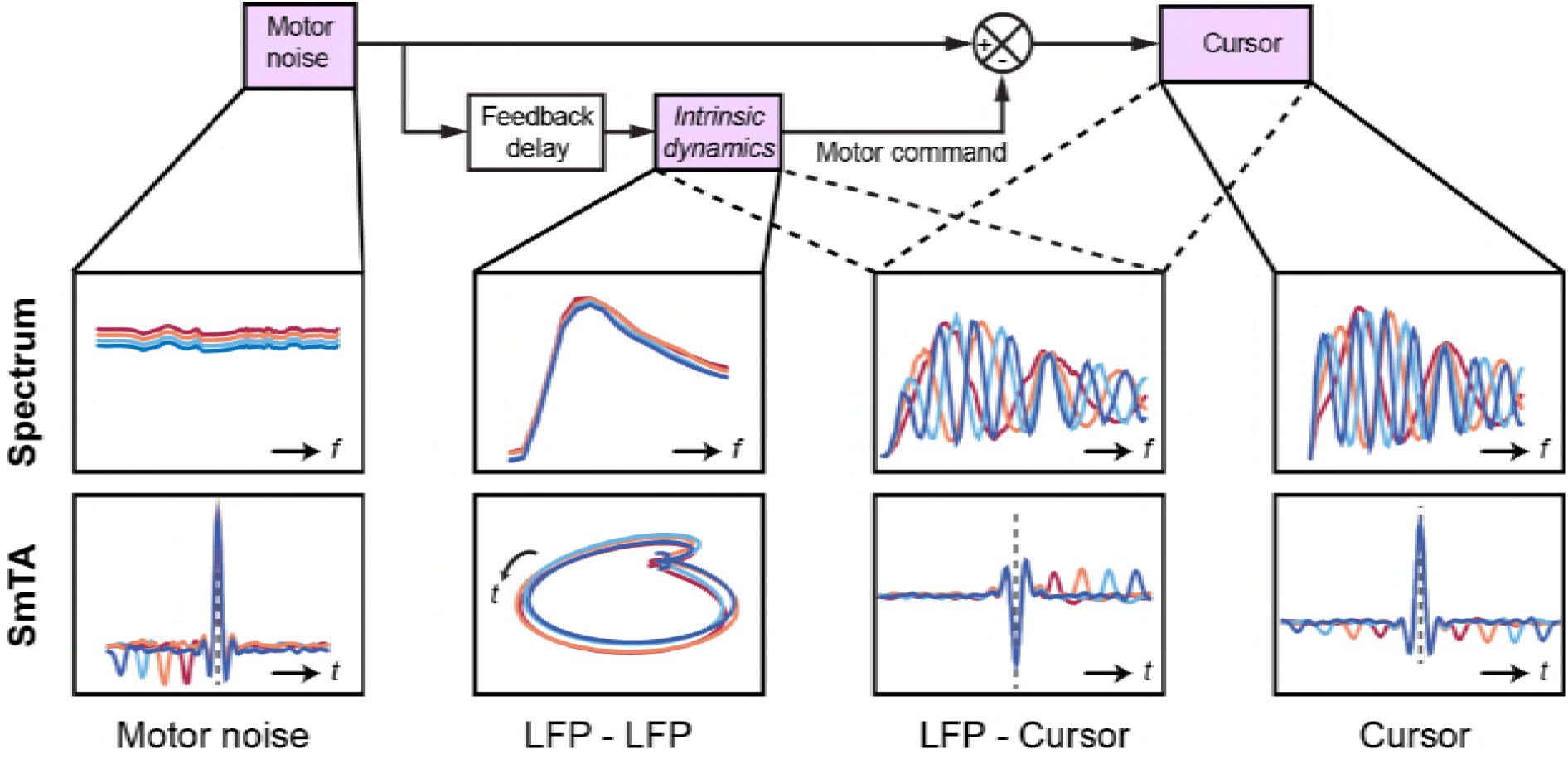
Schematic of delay-dependent and delay-independent relationships in OFC model. The boxes show how the various frequency-domain and submovement-triggered average (SmTA) relationships are explained by the OFC model. *Top row,* from left to right: Broad spectrum motor noise drives intrinsic dynamics resulting in a delay-independent LFP cross-spectral resonance. The delayed motor command is combined with the original motor noise leading to delay-dependent comb filtering evident in LFP-Cursor coherence and Cursor power spectrum. *Bottom row,* from left to right: submovements can arise from a positive noise peak at time-zero, or as a correction to a preceding negative noise trough. Due to intrinsic dynamics, LFPs trace consistent cyclical trajectories locked to submovements. SmTA of LFPs contains potentials associated with noise peak/troughs after feedback delay. SmTA of cursor velocity combines noise with delayed feedback corrections to yield a central submovement flanked by symmetrical troughs.

**Figure S8.**
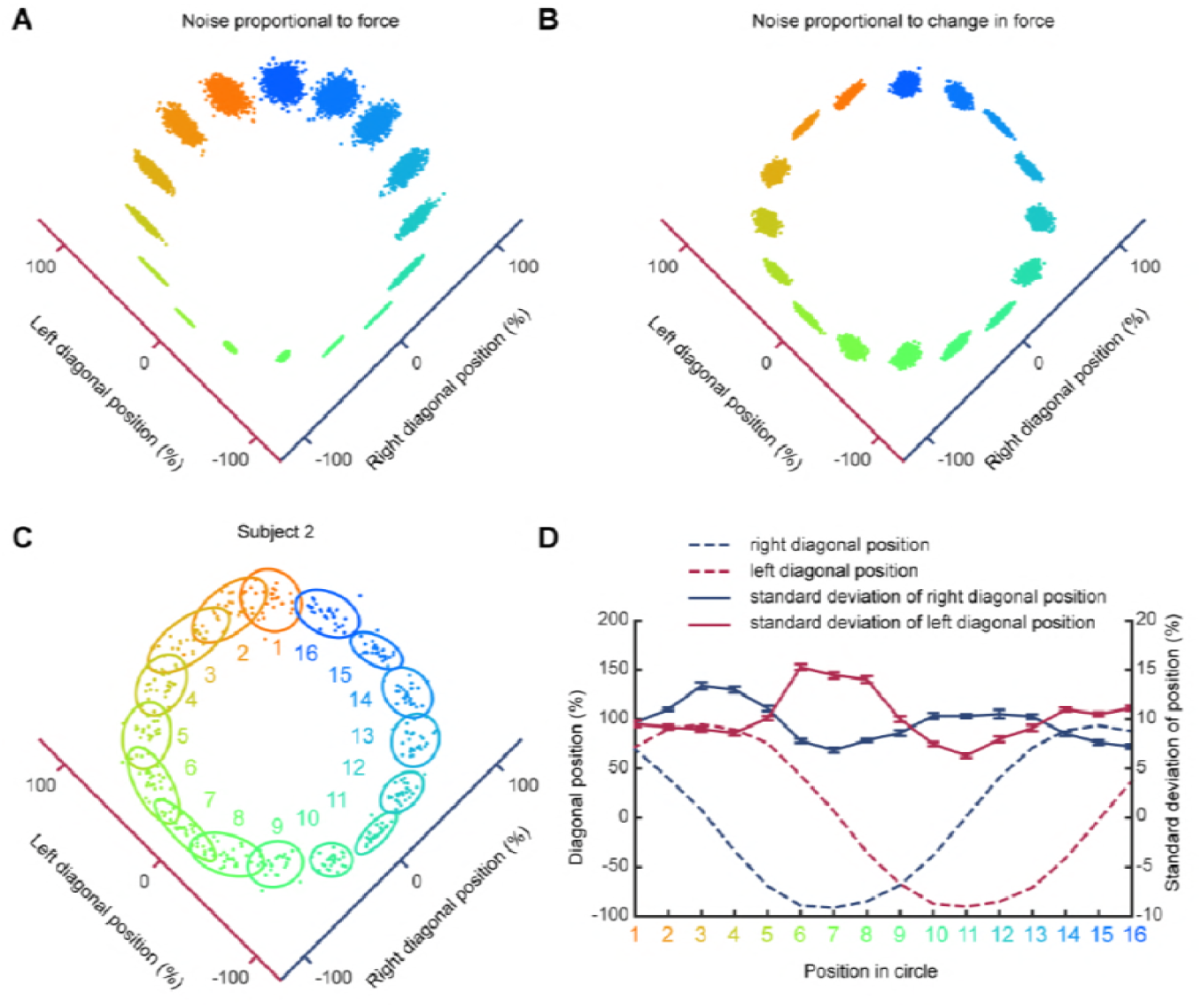
Trajectory variability depends on change in isometric force. (A) Simulated pattern of trial-to-trial variability if motor noise is proportional to absolute force. (B) Simulated pattern of trial-to-trial variability if motor noise is proportional to derivative of force. (C) Variability of a typical subject during counter clockwise tracking. 2D cursor position over multiple trials and associated covariance ellipses are shown for 16 target positions. (D) Average and s.e.m. of standard deviation of force along each finger axis for the 16 target positions. Note that variability is maximal at times of maximal change in associated finger force (*dashed lines*).

